# The hypothalamus as a hub for SARS-CoV-2 brain infection and pathogenesis

**DOI:** 10.1101/2020.06.08.139329

**Authors:** Sreekala Nampoothiri, Florent Sauve, Gaëtan Ternier, Daniela Fernandois, Caio Coelho, Monica Imbernon, Eleonora Deligia, Romain Perbet, Vincent Florent, Marc Baroncini, Florence Pasquier, François Trottein, Claude-Alain Maurage, Virginie Mattot, Paolo Giacobini, S. Rasika, Vincent Prevot

## Abstract

Most patients with COVID-19, caused by severe acute respiratory syndrome coronavirus 2 (SARS-CoV-2), display neurological symptoms, and respiratory failure in certain cases could be of extra-pulmonary origin. Hypothalamic neural circuits play key roles in sex differences, diabetes, hypertension, obesity and aging, all risk factors for severe COVID-19, besides being connected to olfactory/gustative and brainstem cardiorespiratory centers. Here, human brain gene-expression analyses and immunohistochemistry reveal that the hypothalamus and associated regions express angiotensin-converting enzyme 2 and transmembrane proteinase, serine 2, which mediate SARS-CoV-2 cellular entry, in correlation with genes or pathways involved in physiological functions or viral pathogenesis. A post-mortem patient brain shows viral invasion and replication in both the olfactory bulb and the hypothalamus, while animal studies indicate that sex hormones and metabolic diseases influence this susceptibility.

## Main text

SARS-CoV-2 infection is increasingly associated with a wide range of neurological symptoms – headaches, dizziness, nausea, loss of consciousness, seizures, encephalitis etc., as well as anosmia or ageusia – in the majority of patients (*1, 2*). Additionally, many COVID-19 patients with severe disease do not respond well to artificial ventilation or display “silent hypoxia” (*3*), suggesting an extra-pulmonary component to respiratory dysfunction, and cardiorespiratory function and fluid homeostasis are themselves subject to central nervous system (CNS) control. However, despite emerging reports of the post-mortem detection of the virus in the cerebrospinal fluid (CSF) (see for example (*4*)) or brain parenchyma of patients (*5*), little is known about how and under what circumstances SARS-CoV-2 infects the brain.

While the possibility of CNS infection has been largely underestimated due to the common view that angiotensin converting enzyme 2 (ACE2), the only confirmed cellular receptor for SARS-CoV-2 so far (*6*), is absent or expressed only at very low levels in the brain (*7, 8*), and that too exclusively in vascular cells (He et al., bioRxiv 2020; doi: https://doi.org/10.1101/2020.05.11.088500) the majority of these studies have focused on the cerebral cortex, ignoring the fact that other regions such as the hypothalamus, are rich in ACE2 (*9*). Intriguingly, most major risk factors for severe COVID-19 (male sex, age, obesity, hypertension, diabetes); reviewed by (*10, 11*); could be mediated by normal or dysfunctional hypothalamic neural networks that regulate a variety of physiological processes: sexual differentiation and gonadal hormone production, energy homeostasis, fluid homeostasis/osmoregulation and even ageing (*12–14*). The hypothalamus is also directly linked to other parts of the CNS involved in functions affected in COVID-19 patients, including brainstem nuclei that control fluid homeostasis, cardiac function and respiration, as well as regions implicated in the perception or integration of odor and taste (*12, 14–18*).

Here, we investigated the susceptibility of the hypothalamus and related brain regions to SARS-CoV-2 infection by analyzing the expression of ACE2 and the transmembrane proteinase, serine 2 (TMPRSS2), which cleaves the SARS-CoV-2 spike (S) protein, enabling it to be internalized, from existing data from the Allen Human Brain Atlas (AHBA) (*19*). We also used network analysis and pathway enrichment tools to determine which genes and pathways were correlated with this susceptibility, and whether they were similar to differentially expressed genes in the respiratory epithelium of COVID-19 patients (*20*). Next, we verified our hypothesis by immunolabeling for ACE2, TMPRSS2 and other genes of interest in the hypothalamus of control human subjects and the olfactory regions of human embryos, as well as for SARS-CoV-2 S-protein, nucleocapsid (N) protein and double-stranded RNA (dsRNA) in the brain of a patient deceased after severe COVID-19. Finally, we verified whether these cellular and molecular mechanisms of susceptibility could be validated in animal models and whether they were altered by metabolic disorders or variations in sex hormones.

### ACE2 and TMPRSS2 are expressed in the human hypothalamus and connected brain regions

The AHBA contains microarray gene expression data for >62,000 gene probes, including ACE2 and TMPRSS2. We used the atlas to first investigate ACE2 and TMPRSS2 expression in a number of hypothalamic nuclei and connected regions involved in regulating olfaction, gustation or cardiorespiratory function: the insula, amygdala, paraventricular nucleus of the thalamus, pons and myelencephalon (**Fig. 1a**). The frontal lobe, cerebellar cortex and choroid plexus were included for comparison. The OBs and some brainstem regions were not represented in the AHBA. Normalized log2 expression levels of both genes varied widely across and within the brain regions studied (**Fig. 1b**, **Suppl. Table 1, Suppl. Movie**). As expected from previous studies (*21*) (He et al., bioRxiv, 2020; https://doi.org/10.1101/2020.05.11.088500), ACE2 expression levels in the cerebral and cerebellar cortex were relatively low, while the paraventricular hypothalamic nucleus (PVH) displayed the highest ACE2 levels among hypothalamic nuclei, in keeping with its role in fluid homeostasis (*22*). Surprisingly, the choroid plexus displayed extremely high ACE2 levels. TMPRSS2 was present in all areas studied, at levels higher than those of ACE2 on the whole. Relatively high levels of both ACE2 and TMPRSS2 were found in a number of regions, including the PVH and several brainstem areas, indicating that they could be potential targets of SARS-CoV-2.

**Figure 1.**
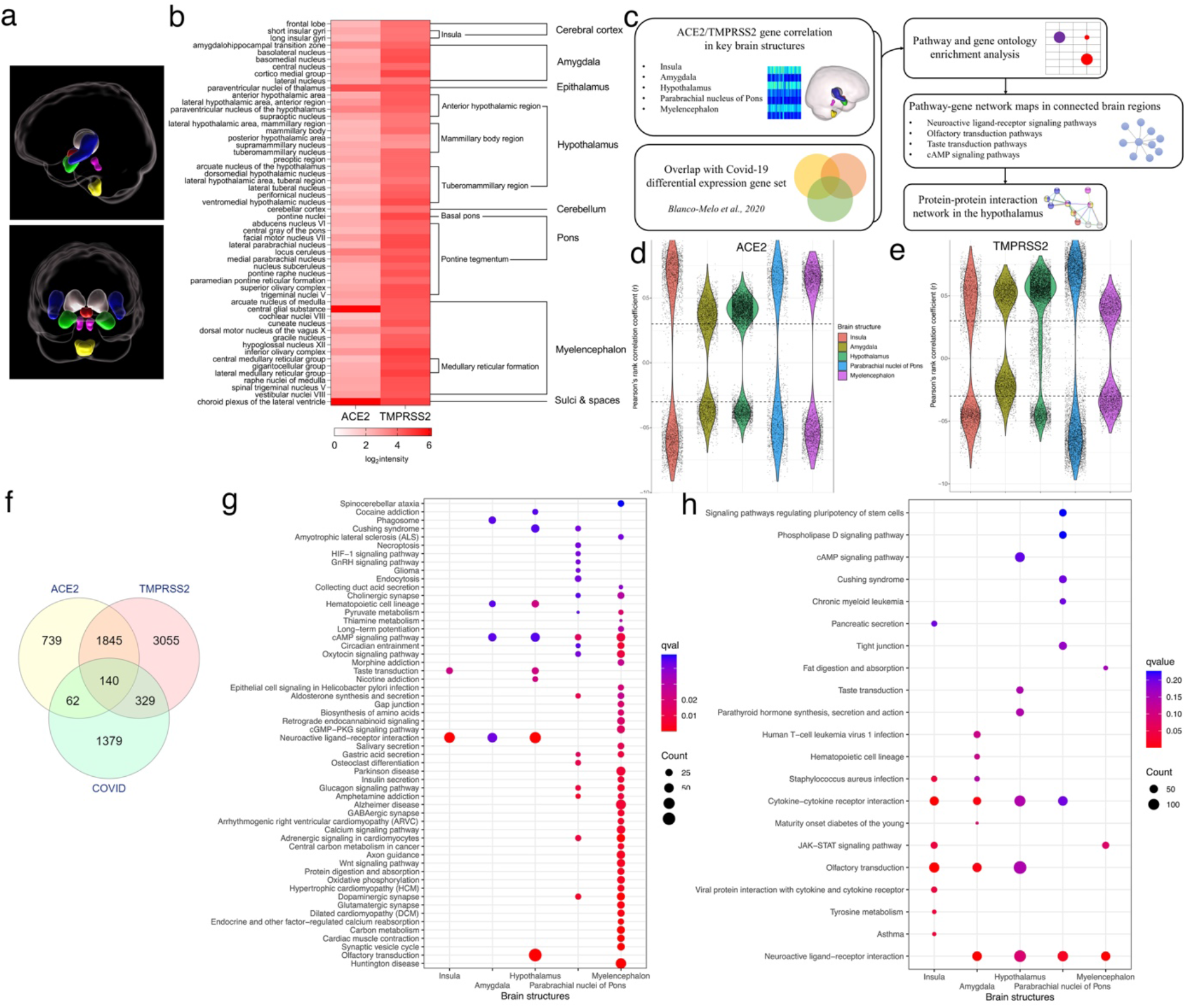
Expression of ACE2, TMPRSS2 and correlated genes and pathways in the human hypothalamus and connected brain regions. (**a**) Schematic of brain regions analyzed in the AHBA in sagittal (top) and frontal (bottom) views. Insula: blue, amygdala: green, thalamus: grey, hypothalamus: red, pons (parabrachial nucleus): pink, myelencephalon: yellow. The frontal lobe of the cerebral cortex, the cerebellar cortex and the choroid plexus were analyzed for comparison but not shown here. (**b**) Heat Map of ACE2 and TMPRSS2 showing log2 expression values in various nuclei or subregions (left) in the different brain regions studied (right). (**c**) Schematic of correlation and interaction analyses of gene-expression data from the AHBA. (**d**,**e**) Violin plots showing number and distribution by Pearson’s rank correlation coefficient (r) of ACE2- and TMPRSS-correlated genes in 5 brain regions. (**f**) Venn diagram showing overlap of hypothalamic ACE2- and/orTMPRSS2-correlated genes (−0.3 < r < 0.3; fdr < 0.25) with COVID-19 patient lung geneset (*20*). (**g,h**) Most significant KEGG pathways yielded by gene enrichment for ACE2-correlated (**a**) and TMPRSS2-correlated genes (**h**) in each of the 5 regions of interest. q value cutoff was set at 0.05 for ACE2 in this figure due to the very large number of significant pathways. Otherwise, q value cutoff was set at an acceptable exploratory level of 0.25. Note that the pathways with the highest number of enriched genes (Count) are in the hypothalamus for both genes queried.

### Genes correlated with ACE2/TMPRSS2 and SARS-CoV-2 infection in key brain structures

We next identified genes whose expression was positively or negatively correlated with ACE2 or TMPRSS2, using their normalized log2intensity gene expression values and the *Find Correlates* option in the AHBA, for 5 regions of interest – the insula, the amygdala, the hypothalamus, the parabrachial nucleus in the pons, and the myelencephalon (**Fig. 1c**). The paraventricular thalamic nucleus, sampled in only three brains in the AHBA, and the pontine tegmentum and basal pons, in which the diversity and number of regions yielded results that were hard to interpret, were excluded from further analysis. However, the parabrachial nucleus of the pons, which was comparable to the diencephalic and telencephalic regions analyzed and key to the functions studied here, was retained. Out of several thousand genes whose expression was correlated with ACE2 or TMRPSS2 in the 5 regions (**Fig. 1d,e**), we identified 2786 with a Pearson’s coefficient (r value) between −0.3 and 0.3, a p value of < 0.01 and a false discovery rate (FDR) of < 0.25 for ACE2, and 5369 for TMPRSS2, in the hypothalamus alone (**Suppl. Table 2**); 1985 were correlated with both ACE2 and TMPRSS2, making them possible candidates of interest for infectious mechanisms or host cell response. In order to draw parallels between SARS-CoV-2 infection of the respiratory epithelium and a putative brain infection, we cross-checked our genes against those differentially expressed in the lungs of COVID-19 patients as compared to healthy individuals (*20*). A total of 140 genes were both correlated with ACE2 and TMPRSS2 expression in the hypothalamus and differentially expressed in COVID-19 patient lungs (**Fig. 1f**, **Suppl. Table 3**). A further 62 and 329 genes, respectively, were correlated with only ACE2 or only TMPRSS2.

### ACE2/TMPRSS2-correlated genes are involved in functional networks of interest

To explore the possible role of ACE2- or TMPRSS2-correlated genes in SARS-CoV-2 pathogenesis, we then performed enrichment analyses on them for KEGG pathways using the *Clusterprofiler* package in R. The KEGG pathway database, which is manually curated, can be used to identify genes known to be implicated in a number of molecular processes and networks. Pathway enrichment and selection yielded a number of important functions and interactions among the ACE2- or TMPRSS2-correlated genes for each brain region analyzed (**Fig. 1g,h**). For ACE2, enrichment in the myelencephalon and pons revealed a very large number of KEGG pathways with few significantly correlated genes, perhaps due to the diverse peripheral inputs they integrate. For this reason, only pathways with a q value of < 0.05 are shown (**Fig. 1g**). In contrast, the insula yielded 2 enriched pathways, and the amygdala 4. The pathways with the most correlated genes, both localized in the hypothalamus, were for “olfactory transduction” (due to a very large number of OR olfactory receptor genes) and “neuroactive ligand-receptor interaction”, in keeping with the variety of neuropeptides and neurotransmitters produced by hypothalamic networks.

KEGG pathway enrichment of TMPRSS2-correlated genes (**Fig. 1h**) revealed very few pathways in the same q value range used for ACE2. We therefore applied an exploratory cutoff of q < 0.25, which is justified for the identification of likely patterns rather than any single significantly correlated pathway. Under these conditions, enrichment revealed a lower number of pathways in the hypothalamus, myelencephalon and pons, but a higher number in the insula and amygdala. Here too, the largest pathways were in the hypothalamus, for “neuroactive ligand-receptor interaction” and “olfactory transduction”, suggesting that these two pathways could be of unusual importance and providing support for our focus on anosmia and neuroendocrine function in SARS-CoV-2 pathogenesis. Other common enriched pathways between ACE2- and TMPRSS2-correlated genes were those for “taste transduction”, “cAMP signaling”, “Cushing syndrome” and “hematopoietic cell lineage”. However the latter two were not enriched in the hypothalamus with regard to TMPRSS2-correlated genes. While this is not in itself an indication that the corresponding genes are not involved in SARS-CoV-2 susceptibility or pathogenesis, we chose not to analyze them further at present.

We additionally performed enrichment for gene ontology (GO) terms (**Suppl. Table 4,5**).

### Functionally connected brain regions express common ACE2- and TMPRSS2-correlated genes and pathways

While, pathway enrichment identifies likely functional links by pinpointing groups of genes that are jointly correlated, the relationship between pathways and individual genes that might connect them or that are similarly correlated in several functionally linked areas could also be informative. We therefore built gene networks based on the pathways identified above for ACE2- (**Suppl. Fig. 1–5**) and TMPRSS2-correlated genes (**Suppl. Fig. 6–10**) in each of the 5 regions. Next, we analyzed the four pathways identified above: “neuroactive ligand-receptor interaction” (**Fig. 2a,b**), “olfactory transduction” (**Fig. 2c,d**), “taste transduction” (**Fig. 2e,f**) and “cAMP signaling” (**Fig. 2g,h**). For each of these networks, a number of ACE2- and/or TMPRSS2-correlated genes were also expressed as part of the same or linked pathways across multiple brain areas, suggesting their involvement in common physiological functions or signaling mechanisms involving ACE2 or TMPRSS2. In addition, both among these common genes and those only present in a single region, we found several, with different functions, whose expression was also altered in the infected lung (*20*), suggesting that they could be involved in susceptibility or response to the virus (**Table 1**). Intriguingly, this list revealed a gene that was highly correlated with both ACE2 and TMPRSS2 in the hypothalamus and also present in other areas: the formyl peptide receptor FPR2 (also known as ALX or FPRL1), a molecule that is closely related to vomeronasal receptors (*23*) but principally acts in immune cells to detect pathogenic peptides and modulate inflammation (*24*).

**Figure 2.**
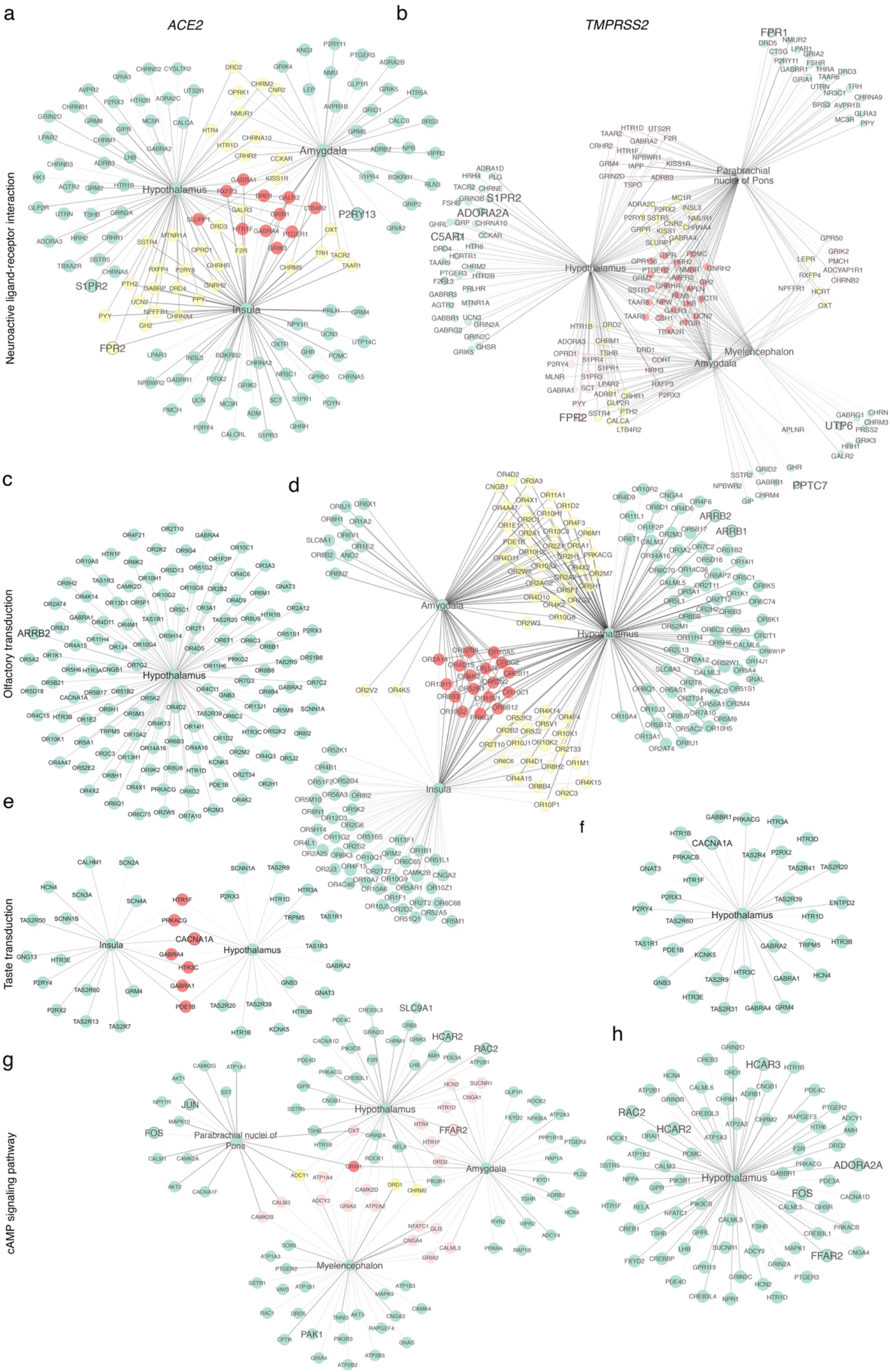
Functionally connected brain regions express common ACE2- and TMPRSS2-correlated genes and pathways. Network of ACE2- and TMPRSS2-correlated genes showing common genes in the four most ubiquitous KEGG pathways in all brain regions where they are enriched. (**a**,**b**) KEGG pathway network for “neuroactive ligand-receptor interaction”. (**c**,**d**) KEGG pathway network for “olfactory transduction“. (**e**,**f**) KEGG pathway network for “taste transduction”. (**g**,**h**) KEGG pathway network for “cAMP signaling pathway”. Node colors: red = genes common to all the regions in which enriched (**a,b,d,e,g**); yellow = genes common to 2 (**a,d**) or 3 (**b,g**) regions in which enriched; pink = genes common to 2 regions in which enriched (**b,g**); green = genes found in only one region. Genes also common to the COVID-19 patient lung dataset are indicated in larger font in all networks.

**Table 1.**
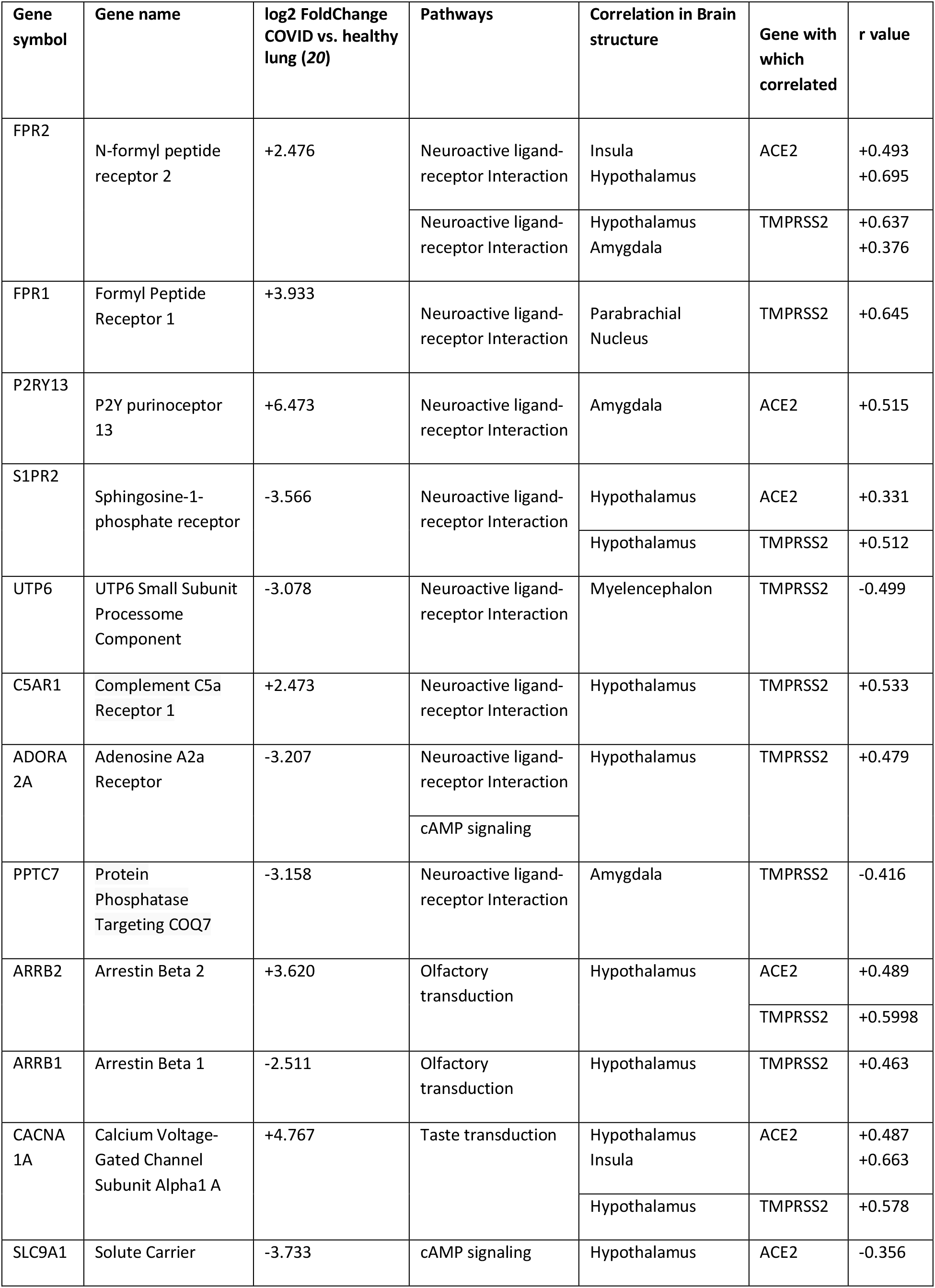
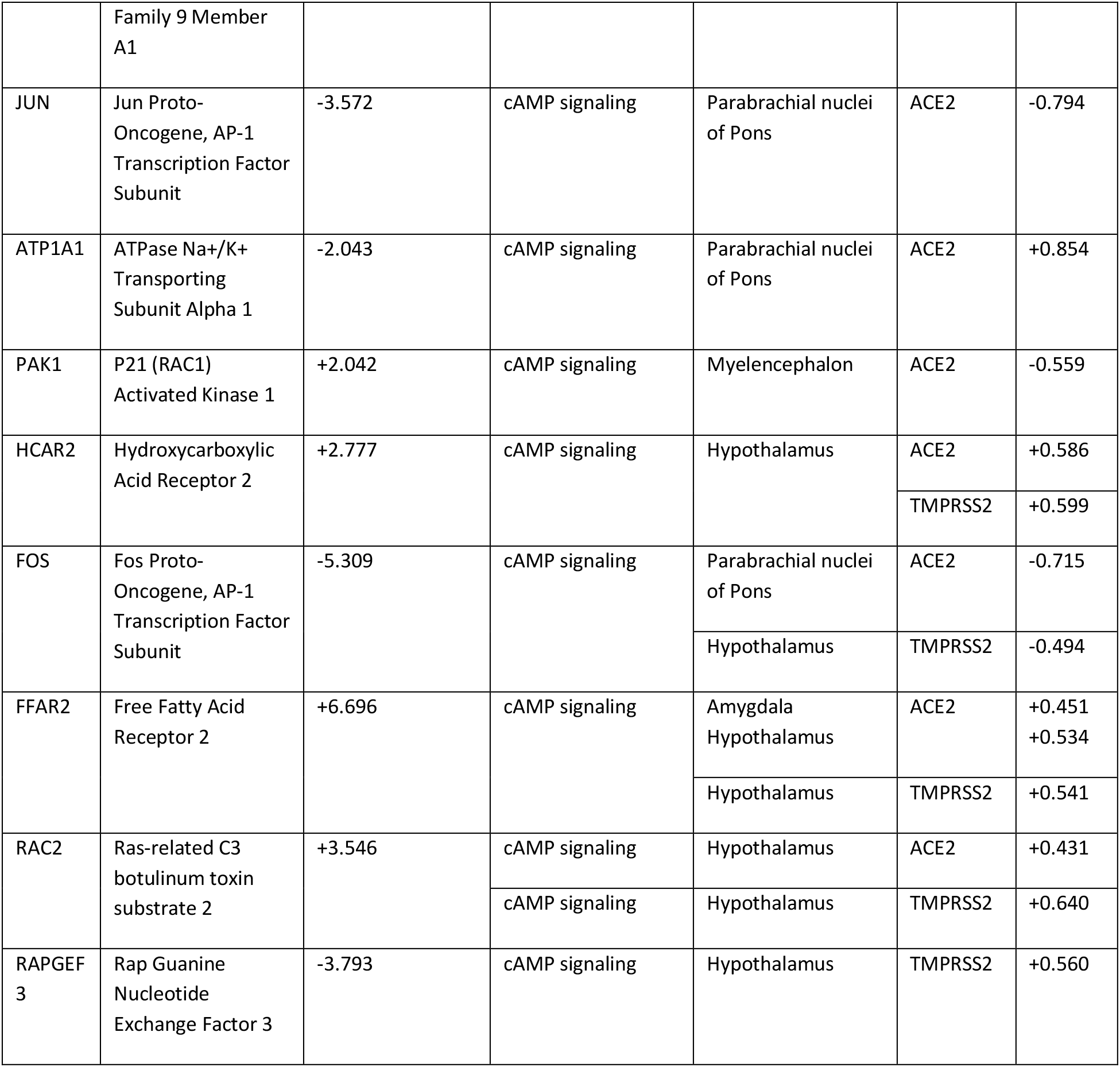
Enriched genes common to multiple regions and the COVID-19 lung dataset. The gene sets identified by our enrichment for the 4 KEGG pathways of interest were cross-checked against the geneset obtained from the COVID-19 lung (*20*), and the common genes annotated below.

### ACE2 and TMPRSS2 are present in human hypothalamic neurons and tanycytes and olfactory sensory neurons

To verify the relevance of our gene expression and network analysis results to human COVID-19 patients, we used available antibodies to perform immunolabeling for ACE2 (**Suppl. Fig. 11**), TMPRSS2 as well as FPR2 on sections of control adult human brains and fetal heads. ACE2 immunoreactivity in the control adult brain was mostly weak, despite tyramide amplification (see Methods). In the PVH parenchyma, ACE2 was present in cells with a neuronal morphology, where it was colocalized with FPR2, as well as a few capillary walls (**Fig. 3a**). Some FPR2-positive fibers or cell bodies, possibly glial, were visible close to the ventricular wall, but were not always colabeled for vimentin (**Fig. 3b**), a marker in the hypothalamus for specialized ependymoglial cells called tanycytes. More ventrally, at the level of the arcuate nucleus of the hypothalamus (ARH) and median eminence (ME), a circumventricular organ (CVO) at which the traditional blood-brain barrier has been replaced by a fenestrated endothelium and a barrier consisting of tanycytes, light ACE2 labeling was seen in a few tanycytic processes radiating from their cell bodies in the ventricular wall (**Fig. 3c**). ACE2 in these processes colocalized with TMPRSS2, suggesting that the machinery for SARS-CoV-2 infection is present in tanycytes. Some capillary walls were also positive for ACE2 as expected, as were some cells with a neuronal morphology (**Fig. 3c**). TMPRSS2 was strongly colocalized with vimentin-positive tanycytic processes in the ME/ARH at both the ventricular wall and the pial surface (**Fig. 3d,e**).

**Figure 3.**
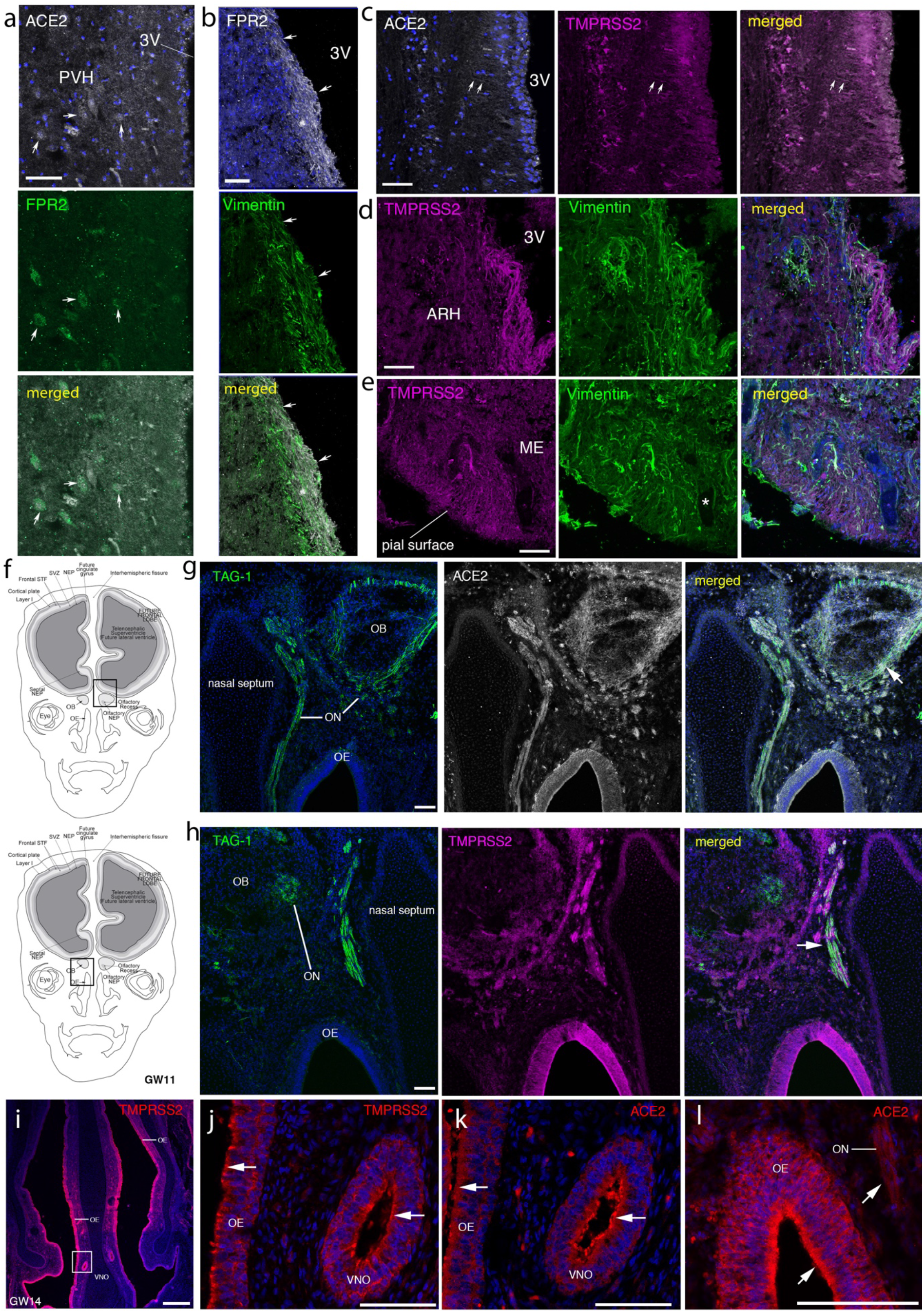
Immunolabeling for ACE2 and TMPRSS2 in the control adult human brain and embryonic human nose and olfactory bulb. (**a**) ACE2 is sparsely present in neurons of the PVH (solid arrows) of a 50 year-old control subject, where it colocalizes with FPR2 (green). (**b**) Closer to the ventricular wall, FPR2 labeling (white) occurs in fibers of unknown origin but rarely colocalizes with the tanycytic marker vimentin (green). (**c**) ACE2 (white) is also present at low levels and colocalized with TMPRSS2 (magenta) in tanycytic processes bordering the 3V (arrows). (**d,e**) TMPRSS2 (magenta) is strongly expressed in vimentin-positive (green) tanycytes in the ARH and ME. (**f**) Schematic diagram of the cross section of an embryonic head at gestational week (GW) 11, showing areas represented. Boxes in (**f**) depict the areas shown in the photomicrographs in (**g,h**). (**g,h**) ACE2 (**g**, white) and TMPRSS2 (**h**, magenta) colocalize (white arrows) with TAG-1 (green), a marker of the axon tracts in the primary olfactory region (olfactory nerve, ON) and of vomeronasal axons, which, at this stage, start projecting to the presumptive olfactory bulbs (OB). (**i**) Low magnification of section of nasal area with inset showing area represented in (**j-l**). (**j-l**) Olfactory and vomeronasal sensory neurons, along with other supporting cells, strongly express ACE2 (**k,l**) or TMPRSS2 (**j**) (both red, white arrows). Blue: DAPI. VNO: presumptive vomeronasal organ. STF: stratified transitional fields; SVZ, subventricular zone; NEP: neuroepithelium. All scale bars = 100 μm.

In order to investigate a putative olfactory route of infection, given the absence of OB data in the AHBA, we performed ACE2 and TMPRSS2 labeling in whole head sections of human fetuses at gestational week (GW) 11 (**Fig. 3f**) or 14 (**Fig. 3i**). Both ACE2 (**Fig. 3g**) and TMPRSS2 (**Fig. 3h**) were strongly colocalized along TAG-1 (or contactin 2, CNTN2)-immunoreactive olfactory/vomeronasal axons, which start targeting the OBs at these developmental stages. Notably, at GW11, all cells in the olfactory epithelium (OE) were positive for ACE2 and TMPRSS2 (**Fig. 3g,h**), and both proteins were strongly expressed in the developing OB (**Fig. 3g,h**). At GW14, when the neuronal layer of the OE and the embryonic vomeronasal organ (VNO) was further developed, strong ACE2 and TMPRSS2 immunolabeling was observed in the apical layer of both the OE and the VNO, in addition to the olfactory nerve (ON), most likely corresponding to olfactory and vomeronasal sensory neurons, respectively (**Fig. 3j,k,l**). FPR2 was not present and is not known to occur in the nose, despite its homology to vomeronasal receptors.

### SARS-CoV-2 attaches to and infects olfactory sensory neurons and hypothalamic median eminence cells in the COVID-19 patient brain

We obtained the brain of a 63 year-old obese male patient who tested positive for SARS-CoV-2 at hospitalization, and who died after 33 days in the ICU after severe respiratory distress, complications and multi-organ failure. We observed strong ACE2 labeling around blood vessels, and both ACE2 and TMPRSS2 labeling in the outer layer of the patient’s OB (**Fig. 4a**, **bottom panels**) but not in a control brain from a 36 year-old overweight male patient negative for SARS-CoV-2 **(Fig. 4a**, **top panels**). Immunolabeling for olfactory marker protein (OMP) confirmed that this layer (ONL) is where the olfactory nerve, consisting of fibers from olfactory sensory neurons, enters the OB (**Fig. 4c**). Interestingly, these fibers were also highly immunopositive for the SARS-CoV-2 S-protein (**Fig. 4a**, **bottom panels**), and immunolabeling revealed dsRNA from replicating viruses in numerous cells bordering the ONL and within the glomerular layer (**Fig. 4c**), indicating viral entry into the brain through olfactory sensory neuronal fibers from the nose. FPR2 was also found at high levels in the ONL in the patient but not the control brain (**Fig. 4b**), suggesting that SARS-CoV-2 infection itself increases the expression of ACE2, TMPRSS2 and FPR2 in olfactory sensory neurons, potentiating infection.

**Figure 4.**
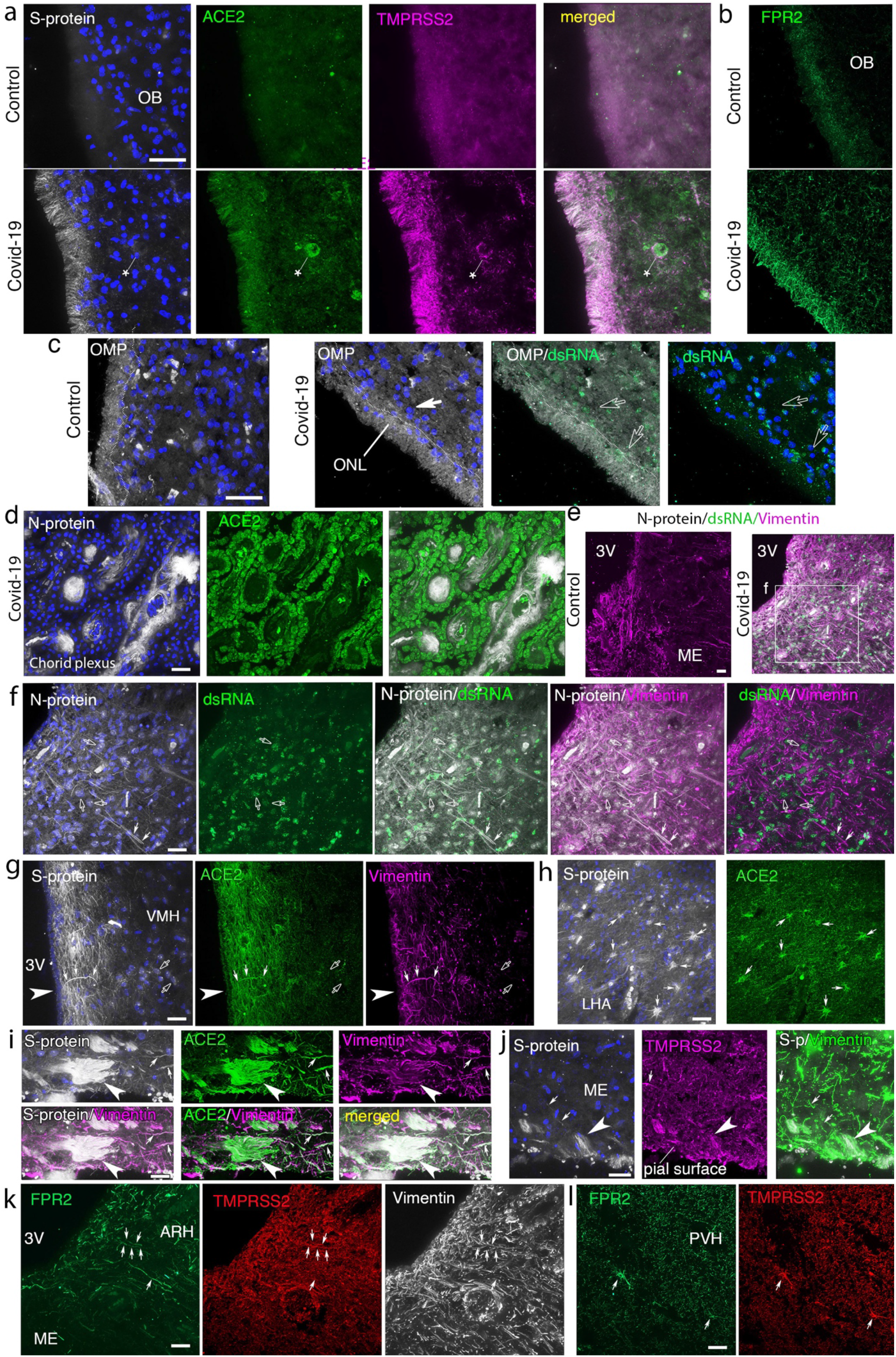
(**a**) Olfactory bulb of a 36-year old control subject (top panels) and a 63 year-old COVID-19 patient (bottom panels) showing heavy viral spike protein (S-protein; white) labeling as well as strong ACE2 (green) and TMPRSS2 (magenta) expression in the outer (olfactory nerve) layer of the olfactory bulb (OB), consisting of the fibers of olfactory sensory neurons in the nose. ACE2 is particularly enriched in the walls of blood vessels (asterisk) as well. Blue – DAPI. (**b**) FPR2 (green) is also highly enriched in the outer layer of the OB in the patient as compared to the control. (**c**) Immunolabeling for the olfactory marker protein (OMP; white), a marker of olfactory sensory neurons, clearly indicates that this layer, delineated by the faint dotted line, is indeed the olfactory nerve layer consisting of the axons of olfactory sensory neurons. Numerous cells of unknown identity containing double-stranded viral RNA (dsRNA; green) can be found close to the border (empty arrowheads) as well as further inside the parenchyma of the glomerular layer, but rarely in the ONL. (**d**) Choroid plexus of the COVID-19 patient showing heavy expression of ACE2 (green) as well as viral nucleocapsid protein (N-protein; white) labeling in blood vessels. (**e**) Immunolabeling for the viral N-protein (white), viral dsRNA (green) and vimentin (magenta) to mark tanycytes, indicating that both viral markers are absent in the hypothalamus of control subjects (left) but heavily expressed in the 63 year-old SARS-CoV-2-infected patient (right). 3V: third ventricle. (**f**) N-protein (white), dsRNA (green) and vimentin (magenta) immunolabeling showing that the N-protein is colocalized with vimentin in a number of tanycytic fibers (white arrows) and present in a number of non-tanycytic cell bodies (empty arrows) throughout the hypothalamic parenchyma near the median eminence. Note that dsRNA is not present in the vimentin-rich tanycytic cell body layer at the edge of the ventricular wall. (**g**) Heavy labeling for the S-protein (white) and ACE2 (green) in fibers lining the wall of the 3V, some capillary walls and some vimentin-positive (magenta) tanycytes (white arrows) at the level of the ventromedial nucleus of the hypothalamus (VMH) in the patient brain; light labeling is also seen in cells with a neuron-like morphology (empty arrows) in the parenchyma. (**h**) Numerous cells with an astrocytic morphology (white arrows) are labeled for both S-protein (white) and ACE2 (green) in the parenchyma of the lateral hypothalamic area (LHA). (**i,j**) Extremely strong labeling for S-protein (white) is seen in the endfeet (white arrowhead) of tanycytes (vimentin; magenta in **i**; green in **j**), which also express ACE2 (green in **i**) and TMPRSS2 (magenta in **j**), at the pial surface of the ME, where tanycytic processes (white arrows) contact the fenestrated endothelium of capillaries in this area. (**k**) In the arcuate nucleus (ARH), FPR2 (green) colocalizes with vimentin (white) and in some cases TMPRSS2 (red), including in a few tanycytic fibers (white arrows). (**l**) FPR2 (green) and TMPRSS2 (red) colocalize in some astrocyte-like cells (white arrows) in the parenchyma of the paraventricular hypothalamic nucleus (PVH), in addition to scattered labeling throughout the parenchyma. Blue – DAPI. Scale bars: 40μm.

ACE2 was strongly expressed in the choroid plexus of the COVID-19 patient, where we also found strong labeling for the highly conserved SARS-CoV nucleocapsid N-protein in what appeared to be blood vessels (**Fig. 4d**).

In the ME/ARH of the COVID-19 patient hypothalamus, abundant labeling for both N-protein and dsRNA could be observed in a large number of cells (**Fig. 4e**, **right panel, f**), indicating robust viral infection and replication. Neither viral marker was seen in the hypothalamus of controls (**Fig. 4e**, **left panel**). However, while diffuse N-protein labeling in the COVID-19 patient was colocalized with vimentin-positive tanycytic fibers in many cases, dsRNA was not present in the cell bodies of tanycytes, which line the wall of the third ventricle (**Fig. 4e**, **right panel, f**). Some parenchymal cells showed high levels of both markers with N-protein aggregation, perhaps indicative of viral assembly. Interestingly, in the ME/ARH, S-protein, present on the outer envelope of assembled viruses, was observed at extremely high levels in the endfeet of vimentin-positive tanycytes at the pial surface of the ME (**Fig. 4i,j**), where it was colocalized both with ACE2 (**Fig. 4i**) and TMPRSS2 (**Fig. 4j**), suggesting that SARS-CoV-2 from the circulation was being internalized by tanycytic endfeet at the level of the fenestrated capillaries, but subsequently transferred to other cell types. In the ventromedial hypothalamus (VMH), however, S-protein was also found in numerous fibers and capillaries close to the ventricular wall, most of which also expressed ACE2; these included some vimentin-positive tanycytes that extended processes into the parenchyma (**Fig. 4g**). In addition, some vimentin-negative cells with a neuron-like morphology in the parenchyma of the VMH (**Fig. 4g**) and cells with an astrocytic morphology in the lateral hypothalamic area (LHA) (**Fig. 4h**) also expressed both ACE2 and S-protein. The widespread and strong expression of ACE2 in the hypothalamus of the SARS-CoV-2 infected patient, unlike the weak expression seen in the control brain, was reminiscent of the increase in ACE2 and TMPRSS2 seen in the ONL of the patient. As for FPR2, at the level of the ME/ARH, it was present both in some vimentin- and TMPRSS2-coexpressing tanycytic processes and in non-vimentin-labeled glial cells that could be microglia, close to the ventricular wall (**Fig. 4k**). More dorsally, it was strongly expressed and colocalized with TMPRSS2 in some cells with an astrocytic morphology in the patient’s PVH (**Fig. 4l**).

### ACE2, TMPRSS2 and FPR2 are expressed in the hamster and mouse hypothalamus and are upregulated by high-fat diet and ovariectomy

To validate our gene and protein expression data in animal models in which to explore viral susceptibility and pathogenesis, we next performed immunofluorescence labeling for ACE2, TMPRSS2 and FPR2 in brain sections from male hamsters, a model susceptible to SARS-CoV-2 infection even without genetic modification due to its higher homology with human ACE2 than mice (*25*). Using the same antibodies to human ACE2 that we used in the human brain sections, we found ACE2 and TMPRSS2 in the processes of a large number of vimentin-positive tanycytes, both in the ME/ARH and more dorsally, spanning the hypothalamic parenchyma from their cell bodies lining the third ventricle to the external surface, where their endfeet contact the capillary bed (**Fig. 5a**). This again underscores the fact that tanycytes could provide an entry mechanism for the virus into the brain.

**Figure 5.**
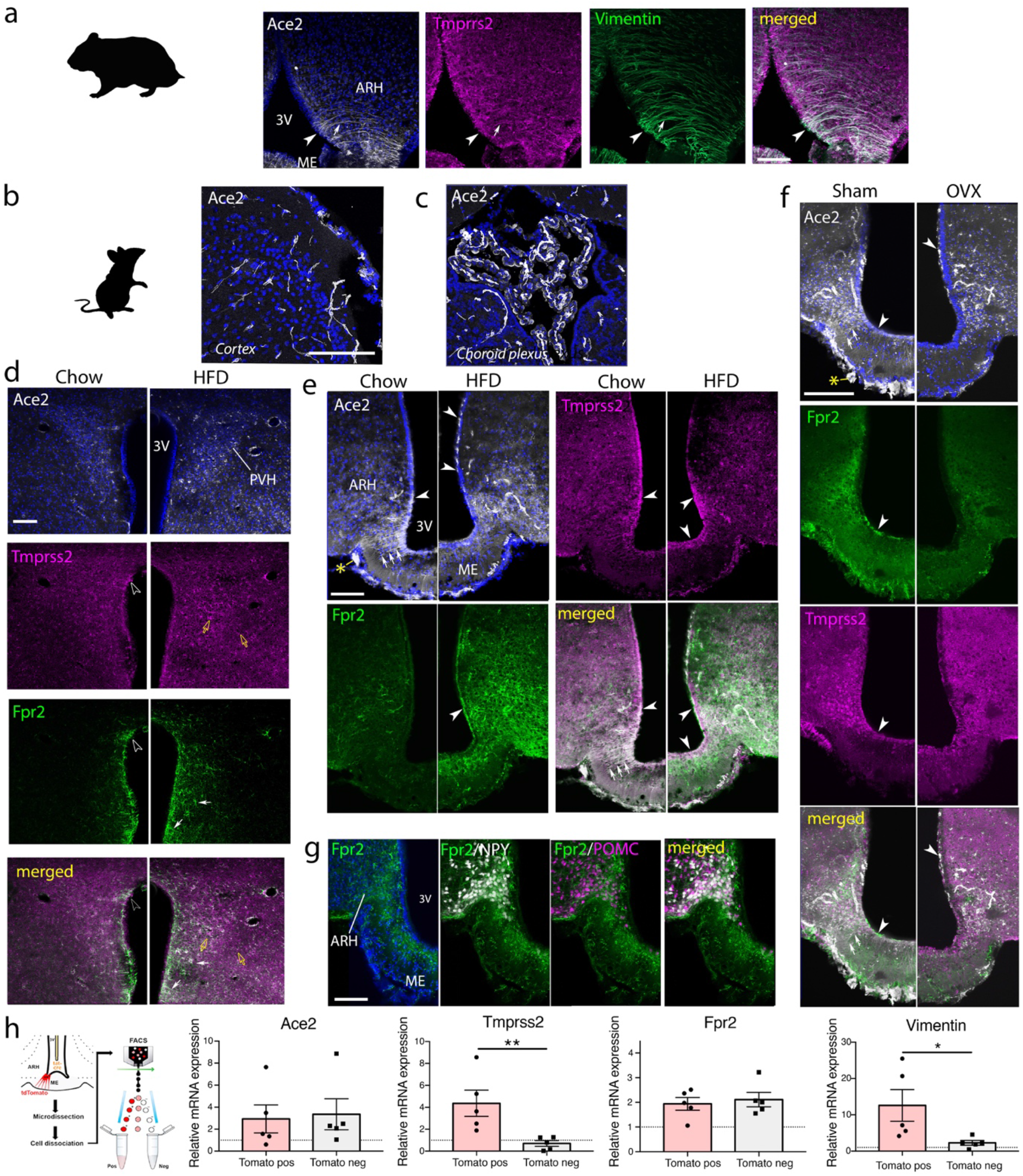
Immunolabeling for ACE2, TMPRSS2 and FPR2 is present and upregulated by high-fat diet in the hypothalamus of mice and hamsters. (**a**) In the ARH and ME of hamsters, ACE2 (white; anti-human ACE2 antibody) and TMPRSS2 (magenta) are colocalized with the tanycytic marker vimentin (green) in both cell bodies bordering the 3V (solid arrowhead) and in processes (solid arrow). (**b,c**) Control regions. In the mouse cerebral cortex, ACE2 (white; anti mouse ACE2 antibody) is limited to the vascular walls (**b**), but it is highly expressed in a polarized pattern in cells of the choroid plexus of the lateral ventricle (**c**). (**d**) ACE2, TMPRSS2 and FPR2 immunolabeling in the paraventricular nucleus of the hypothalamus (PVH), in male mice fed a standard chow diet (left) or a high-fat diet (HFD; right). ACE2 (white) appears to be present in scattered neurons and vascular cells in chow-fed animals, and increases in HFD animals. TMPRSS2 (magenta) is expressed in a few cells with astrocytic morphology at the ventricular wall (empty arrowhead) and scattered microglia-like cells (yellow arrows), with increased expression in the latter in HFD animals. FPR2 (green) is strongly expressed in cells with an astrocytic morphology at the ventricular wall, including in some TMPRSS2-positive cells. It is also present in astrocyte-like cells in the parenchyma near the ventricle. HFD strongly increases the number and area occupied by FPR2-positive cells. 3V: third ventricle. (**e**) ACE2 expression (white) in the hypothalamic arcuate nucleus (ARH) and median eminence (ME) occurs in ME tanycytic cell bodies and processes as well as some blood vessels, and in the pars tuberalis under the vasculature of the ME (asterisk). In HFD animals, ACE2 expression in ME tanycytic cell bodies is reduced and processes are no longer labeled, but labeling appears at the apical pole of tanycytes further up the ventricular wall and increases greatly in capillary walls in the ARH. The labeling in the pars tuberalis is also reduced. TMPRSS2, also present in tanycytic cell bodies and processes, increases slightly with HFD. FPR2, which appears to be in microglia-like cells in the ME and ARH and the cell bodies of tanycytes, increases strongly. (**f**) ACE2 (white), TMPRSS2 (magenta) and FPR2 (green) immunolabeling in the ARH and ME of ovariectomized (OVX) female mice showing changes in ACE2 and TMPRSS2 similar to that seen in male HFD mice. FPR2 labeling, however, appears to be less intense and in different cell types or components from that in males. Asterisk, pars tuberalis. (**g**) FPR2 immunolabeling (green) in tanycytic processes and microglia-like cells in the ARH and EM of transgenic mice expressing NPY-GFP (white) and POMC:Cre; tdTomato mice (magenta). Blue: DAPI. Scale bars = 100 μm. (**h**) FACS-based sorting of tdTomato-positive (pos) and negative (neg) cells from the mediobasal hypothalamus of mice expressing tdTomato exclusively in tanycytes (top panel) to confirm cell-type-specific labeling seen in (**e,f**) for ACE2, TMPRSS2 and FPR2. Vimentin (bottom panel) was used to confirm the purity of FACSed cells. *p<0.05, **p<0.01, Mann-Whitney test.

Despite the usefulness of the hamster as a model for SARS-CoV-2 infection, pathophysiological mechanisms, such as risk factors, are easier to elucidate and validate in mice due to the wealth of literature. We therefore next evaluated the effect on ACE2, TMPRSS2 and FPR2 of a standard or high-fat diet (HFD), which induces obesity and diabetes, and the effect of the gonadal steroid estrogen in mice.

In the brain of mice fed standard chow, we first validated an antibody to mouse ACE2. Immunofluorescence using this antibody in most parts of the adult mouse brain appeared to be limited to cells of the capillary walls – pericytes or endothelial cells (**Fig. 5b**) – as previously described (*21*) (He et al., bioRxiv, 2020; https://doi.org/10.1101/2020.05.11.088500). However, in the choroid plexus, where ACE2 levels were among the highest in the human brain (**Fig. 1b**), ACE2 appeared to be present as an apical cap (**Fig. 5c**), confirming our observations in the patient choroid plexus (**Fig 4d**). In contrast, in the PVH of standard-chow-fed male mice (**Fig. 5d**, **left panels**), ACE2 appeared present in scattered neurons in addition to vascular cells, as seen in the human brain. Intriguingly, however, in the ME of male (**Fig. 5e**, **left panels**) and female mice (**Fig. 5f**, **left panels**), in addition to vascular cells, tanycytic cell bodies and processes showed the most labeling. TMPRSS2 labeling was found predominantly in tanycytes in the ARH and ME, but was also found in astrocyte-like cells at the ventricular wall adjacent to the PVH and some microglia-like cells in the PVH parenchyma (**Fig. 5d,e,f**, **left panels**). The pars tuberalis, a highly vascularized zone under the ME, was also strongly ACE2-positive (**Fig. 5f**, **asterisk top panel**). Surprisingly, FPR2 was highly expressed in cells with an astrocytic morphology along the ventricular wall at the PVH, where it sometimes colocalized with TMPRSS2, and in both tanycytes and microglia-like cells in the ARH and ME (**Fig. 5d,e,g**).

Next, we examined the same three markers in male mice given a HFD for 9 weeks to induce obesity (**Fig. 5d,e**, **right panels**) and in ovariectomized (OVX) female mice (**Fig. 5f**, **right panels**). Strikingly, in the male HFD mice, ACE2 was strongly upregulated in vascular cells in the ARH, while at the same time, ACE2 immunolabeling was reduced in the cell bodies of ME/ARH tanycytes and completely abolished in their processes (**Fig. 5e**, **right panels**). However, apical labeling like that seen in the choroid plexus could be seen in tanycytes in the dorsal ARH, while the pars tuberalis was less strongly labeled (**Fig. 5e**, **right panels**). ACE2 also increased in the PVH with HFD (**Fig. 5d**, **right panels**), while TMPRSS2 increased more moderately with HFD in both the microglia-like cells of the PVH and tanycytic cell bodies in the ME/ARH (**Fig. 5d,e**, **right panels**). Importantly OVX female mice displayed the same changes in ACE2 and TMPRSS2 immunolabeling as male HFD mice, in both tanycytes and vascular wall cells (**Fig. 5f**). On the other hand, the number and spatial extent of FPR2-positive astrocyte-like cells in the PVH and tanycytes in the ARH and ME dramatically increased in HFD mice; in addition, there was an increase in microglia-like FPR2 labeling in the latter, in keeping with HFD-induced inflammation (**Fig. 5d,e**, **right panels**) (*26*). However, these observations were not evident or were even reversed in OVX females (**Fig. 5f**, **right panels**), indicating that metabolic and gonadal hormone-mediated changes may involve pathways or regulatory mechanisms that overlap only partially.

Finally, given the diversity of these results, we used fluorescence-associated cell sorting (FACS) to sort cells from the ME of mice expressing tdTomato selectively in tanycytes (*27*) (**Fig. 5h**), and performed RTqPCR for ACE2, TMPRSS2, FPR2 and the tanycytic marker vimentin. While ACE2 and FPR2 were present in both tanycytes and non-tanycytic cells, as indicated by immunolabeling (**Fig. 5e,f**), TMPRSS2 was expressed almost exclusively in tanycytes in this region. Together, these results suggest that ACE2 and FPR2 expression form part of an inflammation-modulating pathway in vulnerable hypothalamic regions, through which risk factors such as metabolic diseases and male sex (or the absence of female sex) could increase susceptibility to and outcome of SARS-CoV-2 infection.

## Discussion

Six months and several thousand papers and preprints after the beginning of the pandemic, if there is one thing we have learnt about SARS-CoV-2, it is that almost every assumption that has been made about the virus has been wrong. Although viral pneumonia, complicated by the “cytokine storm” and a prothrombotic state (*28–31*), is still the principal symptom in severely ill patients, other tissues, notably the gut, are also directly susceptible to infection (*32*). While our study may seem at first sight to resemble the parable of the blind men and the elephant, we consider the possibility that SARS-CoV-2 infiltrates the brain, and specifically the hypothalamus, with functional consequences to disease progression and outcome, to be more in the nature of the elephant in the room. Our current observations of SARS-CoV-2 S- and N-proteins as well as dsRNA in cells of the hypothalamic ME lay to rest any doubt in this regard.

In the hypothalamus, the virus appears to adopt a remarkable infection method involving several cell types, with the spike protein binding to capillary walls and the endfeet of ME tanycytes where they contact the fenestrated endothelium typical of CVOs, viral particles entering the tanycytes and producing more nucleocapsid components there, and the viral RNA transfecting other hypothalamic cells in which it replicates, rather than in tanycytes themselves. ACE2, which we show to be present in tanycytes and highly enriched in the capillary bed of the median eminence, and TMPRSS2, highly expressed by tanycytes themselves, could allow the first step in this infection process. Tanycytes are closely connected to each other in the ARH by gap junctions(*33*) and control the electrical activity of neurons of the ARH and ME (*34*), as well as serving as transcytotic shuttles for blood-borne signals into the CSF (*35*); the machinery underlying these functions – vesicular trafficking, cell-cell communication, membrane fusion etc. – could equally be hijacked to allow virus propagation and dissemination in the brain, including through the CSF. Indeed, the highly related SARS-CoV was also observed by several investigators in post-mortem human brain tissue, including the hypothalamus (*36*). Furthermore, altered labeling for several hormones in the pituitary (*37*) and long-term neuroendocrine deficits in some SARS survivors (*38*) strongly support the notion of the hypothalamus as a target of viral infection.

A frequent criticism of the hematogenous route hypothesis of brain infection has been the fact that the virus is rarely detected by PCR in the blood of patients. However, a closer look at the literature reveals that the virus is not only detected in a variable percentage of infected individuals, but that the chances of detecting it are higher in severely ill patients (for example, see (*39*)). Once in the bloodstream, the virus could also access other parts of the brain through a leaky endothelial barrier, or other CVOs such as the organum vasculosum of the lamina terminalis, the subfornical organ or the area postrema (*35*), all of which play key roles in either the risk factors or the physiological functions targeted by SARS-CoV-2. While the AHBA does not allow gene expression in these CVOs to be analyzed, further immunolabeling experiments may provide evidence for viral targeting.

On the other hand, the idea that SARS-CoV-2 could infect the brain through peripheral nerves, and specifically an olfactory route, has been raised by several authors, both in light of the anosmia observed in COVID-19 patients (*40, 41*) and from experimental animal studies on SARS-CoV (*42, 43*). In the latter, SARS-CoV administered intranasally was found in the hypothalamus and brainstem within days, a determining factor for mortality (*43*). However, although changes to the OB of a COVID-19 patient with anosmia have been noted (*44*), a more recent post-mortem neuropathological study has revealed no anatomical, histological or immunohistochemical changes in the OB or brain in a series of 18 patients, although the “inferior frontal lobe with olfactory tract/bulbs” was positive for the virus by PCR in one case (*45*). In another preprint, while Neuropilin-1-positive olfactory neuronal progenitors in the OE were labeled for S-protein, in the OB itself, only cells of the vascular wall were found to be labeled (Cantuti-Castelvetri et al., https://doi.org/10.1101/2020.06.07.137802), in keeping with the interferon response (*7*) and a presumed microglial barrier in this region (*46*). This is contradicted by our observations not only of heavy S-protein labeling in the ONL, i.e. in mature olfactory sensory neuronal axons, but of numerous dsRNA-positive cells in the glomerular layer in our patient, confirming that the virus indeed infects putative neuronal cells of the OB. Furthermore, ACE2 and TMPRSS2 appear to be upregulated by the virus itself in the adult ONL, as postulated by other authors (*7*), while fetal olfactory and vomeronasal sensory neurons express both ACE2 and TMPRSS2 at high levels. It should be remembered here that the ON and vomeronasal nerve serve as a scaffold to guide gonadotropin-releasing hormone (GnRH) neurons from the nose to the hypothalamus during development (*47*), a process for which Neuropilin-1 is essential (*48, 49*), and that a continuum of GnRH neurons persists in adulthood, directly connecting olfactory areas to the hypothalamus. Additionally, the theoretical ability of the S-protein to bind to alternative receptors (*50*) or be cleaved by other proteases (*51*) makes it possible that molecules other than ACE2 and TMPRSS2 also mediate viral entry, expanding the range of susceptible cells, and the putative *trans*-activation of the virus could obviate the need for the coexpression of the two molecules on the same host cell (*52*).

Regardless of its point of entry, once in the brain, the virus could infect a number of secondary regions through transsynaptic or other mechanisms. However, given the high incidence of anosmia/ageusia, the likelihood of central respiratory dysfunction, and the relationship between hypothalamic neural circuits and risk factors for severe COVID-19, we focused our gene expression analysis on regions implicated in these functions. Surprisingly, among the four most ubiquitous enriched pathways for ACE2 and/or TMPRSS2-correlated genes, we found olfactory and taste transduction. Both OR and TAS genes occur in “ectopic” locations within and outside the brain, and could play roles other than odor or taste perception, such as the sensing of peripheral metabolites by hypothalamic tanycytes (*53–55*) or the mediation of viral infection or the host response (*56, 57*), and could be over- or under-expressed in disorders like obesity or diabetes (*58*).

Our network analysis and immunolabeling studies also unexpectedly revealed the inflammatory mediator FPR2 as possibly playing a key role in brain infection through ACE2. While FPR2 can mediate certain anti-inflammatory effects, it appears involved in the replication of dsRNA, an important step in the propagation of RNA viruses (*59, 60*), and the hypothalamus exhibits inflammation in case of obesity (*61*). Additionally, the deletion of FPR2 in mice alleviates HFD-induced obesity and insulin resistance by suppressing pro-inflammatory mechanisms in the periphery (*62*), suggesting that FPR2 could potentially exacerbate the effects of SARS-CoV-2 infection in patients at risk. Paradoxically, we have recently shown that anorexia induced by systemic proinflammatory cytokines is also mediated by hypothalamic tanycytes, through the IKK subunit Nemo or IKBKG (*63*), which is differentially expressed in the lung of COVID-19 patients (*20*). Additionally, the changes we observed in hypothalamic ACE2 expression, and especially in ME/ARH tanycytes, in ovariectomized mice were strikingly similar to those in HFD mice, suggesting that the lack of estrogens and metabolic imbalances trigger at least partially overlapping molecular mechanisms. This is of interest as various ME/ARH neuronal populations that control feeding and energy expenditure in response to peripheral metabolic signals express both neuronal nitric oxide synthase (nNOS) (*64*), itself found to inhibit the replication of SARS-CoV (*65*), and the estrogen receptor ERα (*64*), linking sex, metabolism and viral susceptibility or inflammation. Again, this is only a part of the picture, as both ACE2 and TMPRSS2 are also themselves responsive to gonadal hormones – estrogens and androgens respectively (*66, 67*), and the role of FPR2 needs to be explored further.

There are no doubt several other networks that are involved both in hypothalamic functions and in viral susceptibility or pathogenicity. As an intellectual exercise, we performed enrichment for GO terms on the 140 genes that were both correlated with ACE2 and TMPRSS2 in the hypothalamus, and differentially expressed in the COVID-19 patient lung. We then used STRING to build a protein-protein interaction (PPI) map using 5 GO terms for biological processes (out of 115) that we felt covered the largest range of functions of interest to a potential SARS-CoV-2 infection of olfactory or neuroendocrine circuits: “regulation of secretion by cell”, “vesicle-mediated transport”, “immune system process”, “cell surface receptor signaling pathway” and “second-messenger-mediated signaling”. The resulting PPI network (p value = 2.58e-09) allowed us to see potential functional relationships between our highlighted proteins, including several that we were unaware of, and identify proteins that appeared to play key roles at the intersection of several functions (**Fig. 6**). This was particularly the case with FPR2, which has previously been noted in hypothalamic microglia (*68*), and which, in addition to binding viral peptides (see for example (*60, 69*)), could play a role in neurodegenerative disorders, and mediate either pro-inflammatory or anti-inflammatory mechanisms (reviewed in (*70–72*)).

**Figure 6.**
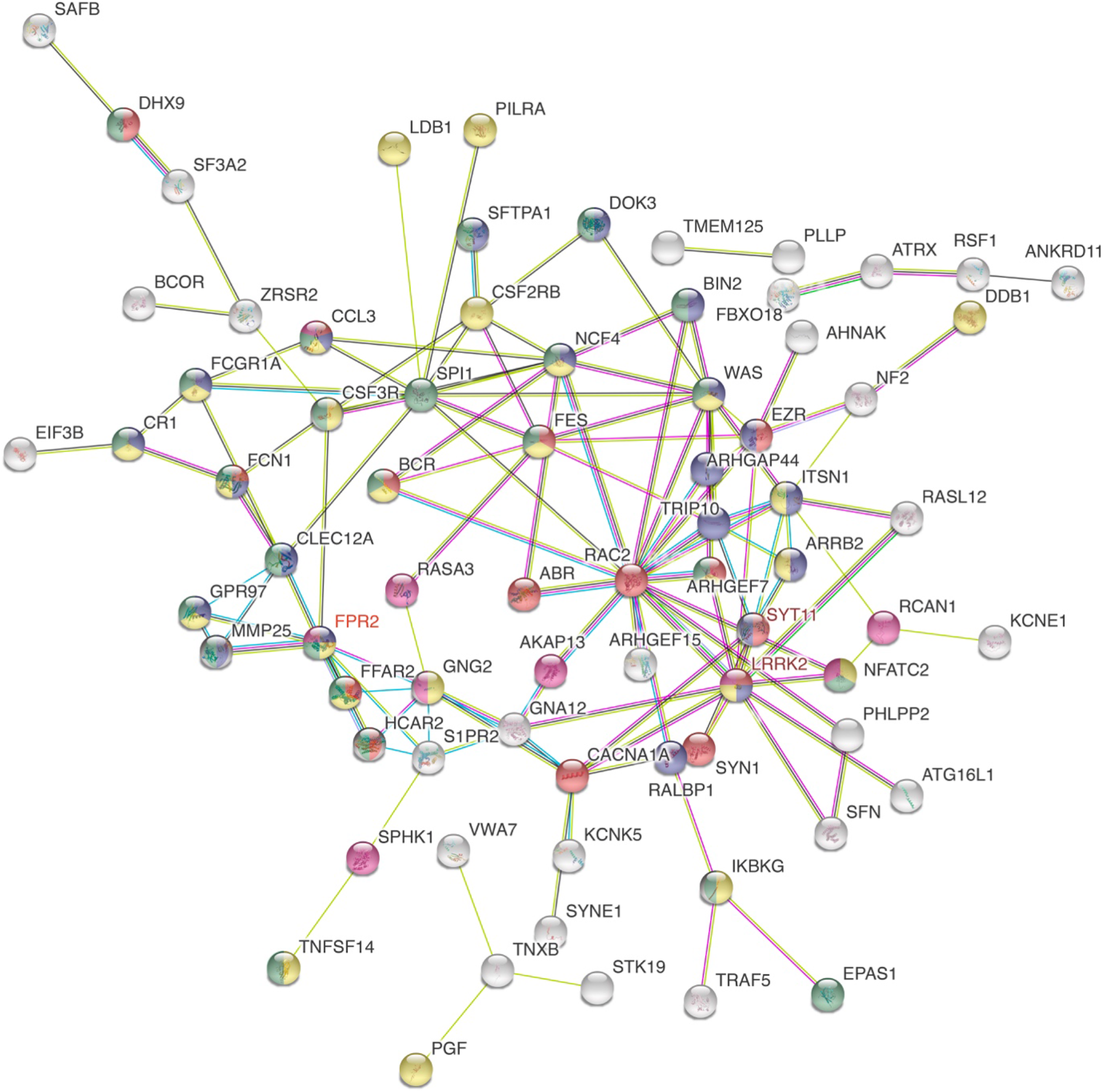
Sample protein-protein interaction map of GO terms of importance to hypothalamic function and viral pathogenesis. Common genes between ACE2-/TMPRSS2-correlated genes in the hypothalamus and the COVID-19 lung differentially expressed geneset were enriched for GO terms for biological processes and queried with STRING to obtain functional protein-protein interaction (PPI) networks using 5 diverse terms of interest: “regulation of secretion by cell” (red), “vesicle-mediated transport” (blue), “immune system process” (green), “cell surface receptor signaling pathway” (yellow), “second-messenger-mediated signaling” (violet). White proteins are connectors not themselves present in our genesets. Empty nodes are those whose structure is not known. Proteins with no interactions highlighted by our analysis were removed to simplify the figure. Note that FPR2 is among the 4 proteins that appear under the most GO terms selected (4 out of 5), along with CCL3, FCN1 and LRRK2. Edge color code for associations: turquoise – known interactions from curated databases; magenta – experimentally determined known interactions; green – predicted interaction based on gene neighborhood; red – predicted interaction based on gene fusions; dark blue – predicted interactions based on gene co-occurrence; yellow – textmining; black –coexpression; pale blue – protein homology. Edges indicate that proteins jointly contribute to a shared function, without necessarily binding to each other.

To summarize, our work provides proof that the brain not only possesses the cellular and molecular machinery necessary to be infected, but that the hypothalamus, which harbors neural circuits regulating a number of risk factors for severe COVID-19 in addition to being linked to olfactory and brainstem cardiorespiratory centers, could be a preferred port of entry and target for the virus.

## Methods

Methods, including statements of data availability and associated accession codes and references are available on line.

## Acknowledgments

The authors are grateful to Dr. Nihal Altan-Bonnet for her valuable insights on the biology of SARS-CoV-2. They would also like to thank Meryem Tardivel (Confocal microscopy) and Nathalie Jouy (FACS) from the BioImaging Center of Lille (BiCeL), and Julien Devassine (animal core facility) of the UMS2014-US41 for their expert technical support.

## Author contributions

S.R. and V.P. designed the study, analyzed data, prepared the figures, and wrote the manuscript. S.N. designed and performed the bioinformatics analyses, and was involved in all aspects of study design, interpretation of results, and manuscript preparation; F.S., G.T., D.F., C.C., M.I. and E.D. prepared tissues and performed the immunofluorescence and RTqPCR analyses; R.P., V.F., M.B., F.P., F.T., C.A.-M, V.M. and P.G. were involved in the study design, interpretation of the results, and preparation of the manuscript.

## Supplementary Materials

Materials and Methods

Figures S1-S10

Tables S1-S5

Movie 1

References (*91-99*)

## Supplementary Materials

## Materials and Methods

### Gene expression analysis for ACE2, TMPRSS2 and correlated genes in the brain

#### Differential distribution of ACE2 and TMPRSS2 across brain regions

The Allen Human Brain Atlas (AHBA) (*19*) comprises 3702 anatomical brain regions from six donors (five males and one female, mean age 42, range 24–57 years). Normalized gene expression values of ACE2 and TMPRSS2 were retrieved from the AHBA for nuclei of the hypothalamus, insula, amygdala, paraventricular nucleus of the thalamus, pons and myelencephalon. The frontal lobe and cerebellar cortex as well as the choroid plexus were included for comparison. The probes were mapped to ACE2 and TMPRSS2 genes using the *collapseRows* function of WGCNA_1.69-81 (*73*) in R ver3.6.3. Probe *CUST_16267_PI416261804* for ACE2 and probe *A_23_P29067* for TMPRSS2 were selected based on maximum variance across all samples (maxRowVariance of WGCNA package). The data were concatenated across all donors for the specified brain structures. To account for missing ACE2 and TMPRSS2 expression values for the brain structures excluded for various technical reasons such as damaged or missing tissue or low-quality RNA (Allen Human Brain Atlas, technical white paper: microarray survey), nonparametric missing value imputation was performed using R-package missForest (*74*). Median expression values of ACE2 and TMPRSS2 across all donors were presented in the form of a heatmap using GraphPad Prism 8.

#### ACE2/TMPRSS2-correlated genes in key brain structures

For the selected ACE2 and TMPRSS2 probes, the “positive” and “anti-correlated” genes with their normalized log2intensity gene expression values were retrieved using “*Find Correlates” search utility in* the AHBA (with a cut-off of −0.3 > r > 0.3 as in the AHBA, where r is Pearson’s rank correlation coefficient) for the brain structures of interest – insula, amygdala, hypothalamus, parabrachial nuclei of pons and myelencephalon - sampled in all human donors. Probes with zero entrez-id were filtered out, and if multiple probes represented a single gene, the one with the lowest adjusted p value was retained. The expression values of the retrieved gene list were used as vector elements to recompute Pearson’s correlation coefficients (r) of all gene pairs using the rcorr function of R-package Hmisc (https://cran.r-project.org/web/packages/Hmisc/index.html). The rcorr function returns a correlation matrix with the r values and the corresponding asymptotic p value based on t distribution. The function p.adjust and the method ‘fdr’ were applied to control for false discovery rate (fdr). ACE2- and TMPRSS2-correlated genes were extracted from the correlation matrix and filtered by (i) setting a threshold of −0.3 > r > 0.3 and fdr < 0.25, and (ii) selecting the correlations with the lowest fdr value for probes mapped to multiple genes. According to the GSEA User Guide, an FDR of <0.25 is reasonable in an exploratory setting, where the aim is to find candidate hypotheses, and where a more stringent FDR cutoff may lead to potentially significant results being overlooked because of the lack of coherence in most expression datasets (see also (*75*)).

#### COVID-19 lung RNA sequencing dataset

Raw read counts from COVID-19 infected and uninfected human lung (n=2) RNA-seq dataset were collected from <u>GSE147507</u> (*20*) followed by differential expression analysis using DESeq2 (*76*). The differentially expressed genes were checked for the overlapping gene count between the COVID-19 dataset and the ACE2- and TMPRSS2-correlated gene sets.

#### KEGG pathway and gene ontology enrichment analyses

KEGG pathway and gene ontology (GO) enrichment analyses of ACE2- and TMPRSS2-correlated genes was performed using *Clusterprofiler* package in R (*77*), a widely used R package for enrichment analyses. enrichGO and enrichKEGG functions based on hypergeometric distributions were applied to perform the enrichment test for the ACE2 and TMPRSS2 correlated gene sets. q value cut-off (for fdr) was consistently maintained at 0.25. The enriched pathways for ACE2 and TMPRSS2 correlated genes for the analyzed brain structures were compiled together and visualized using the ggplot2 package of R. The enriched pathway-gene networks were generated using the cnetplot function of the *Clusterprofiler* package.

#### Gene-network maps connecting brain regions

The gene network maps connecting brain regions with shared pathways such as neuroactive ligand receptor interaction, olfactory transduction, taste transduction and cAMP signaling enriched for ACE2- and TMPRSS2-correlated genes were generated by curating genes associated with each pathway and projecting them in the form of networks using Cytoscape v3.8.0. Furthermore, these genes were checked for overlap with the COVID-19 differential gene expression dataset.

#### Functional protein-protein interaction network in the hypothalamus

Common genes between the COVID-19 differentially expressed geneset and ACE2-/TMPRSS2-correlated genes in the hypothalamus, were queried in the STRING database (https://string-db.org/) to obtain functional protein-protein (PPI) interaction networks, and the associated GO terms enriched for biological processes.

### Collection and processing of human tissues

#### Adult human post-mortem brains

Tissues were obtained in accordance with French bylaws (Good Practice Concerning the Conservation, Transformation and Transportation of Human Tissue to be Used Therapeutically, published on December 29, 1998). Permission to use human tissues was obtained from the French Agency for Biomedical Research (Agence de la Biomedecine,Saint-Denis la Plaine, France, protocol no. PFS16-002) and the Lille Neurobiobank. Studies were undertaken on the of the following subjects: (i) a 63 year-old male patient with obesity and hypothyroidism, who died of multiple complications and organ failure consequent to severe SARS-CoV-2 infection, confirmed by PCR on nasal swabs upon admission to hospital for severe respiratory distress on Day 8 following the appearance of symptoms; (ii) a 36 year-old male patient, overweight, smoker, who died of suspected myocardial infarction after being admitted to the emergency room. PCR for SARS-CoV-2 was negative in all tissues tested (brain, lungs, tracheal and rectal swabs); (iii) a control 50 year old man with no brain disease at the time of death, which occurred before the COVID-19 pandemic.

Dissected blocks of the adult brain containing the olfactory bulb, choroid plexus and/or hypothalamus were fixed by immersion in 4% paraformaldehyde in 0.1M phosphate buffer, pH 7.4 at 4°C for 2 weeks. The tissues were cryoprotected in 30% sucrose/PBS at 4°C overnight, embedded in Tissue-Tek OCT compound (Sakura Finetek), frozen in dry ice and stored at −80°C until sectioning. For human hypothalamus immunolabeling, a citrate-buffer antigen retrieval step, 10mM Citrate in TBS-Triton 0.1% pH 6 for 30 min at 70°C, was performed on 20µm sections. After 3 washes of 5 minutes with TBS-Triton 0.1%, sections were blocked in incubation solution (10% normal donkey serum, 1mg/ml BSA in TBS-Triton 0.1% pH 7,4) for 1 hour. Blocking was followed with primary antibody incubation (see Antibody table) in incubation solution for 48h at 4°C. Primary antibodies were then rinsed out, before incubation in fluorophore-coupled secondary antibodies or, in case of amplified immunolabeling, biotinylated secondary antibodies for 1h in TBS-Triton 0.1% at room temperature. For classic immunohistochemistry, secondary antibodies were washed and sections counterstained with DAPI (D9542, Sigma). For amplified immunohistochemistry, after secondary antibodies were rinsed, sections were incubated with VECTASTAIN® Elite ABC-HRP kit (PK-6100, Vector laboratories) following manufacturer’s instructions. Sections were then incubated with biotinyl-tyramide reagent (SAT700001EA, Perkin Elmer) following manufacturer’s recommendations, washed and incubated with fluorophore-coupled streptavidin (1/500 dilution in TBS-Triton 0.1%) before counterstaining with DAPI. Finally, the sections were incubated with Autofluorescence Eliminator Reagent (2160, Millipore) following manufacturer’s instructions and mounted with Fluoromount™ (F4680, Sigma). Immunolabeling in the human brain using the two antibodies to human ACE2 (R&D Systems, with tyramide amplification, and Abcam, without amplification), labeled similar cells (**Suppl. Fig. 11**).

#### Human fetuses

Tissues were obtained in accordance with French bylaws (Good Practice Concerning the Conservation, Transformation, and Transportation of Human Tissue to Be Used Therapeutically, published on December 29, 1998). The studies on human fetal tissue were approved by the French agency for biomedical research (Agence de la Biomédecine, Saint-Denis la Plaine, France, protocol n: PFS16–002). Non-pathological human fetuses (11 and 14 gestational weeks (GW), n = 1 per developmental stage) were obtained from voluntarily terminated pregnancies after written informed consent was obtained from the parents (Gynecology Department, Jeanne de Flandre Hospital, Lille, France). Fetuses were fixed by immersion in 4% PFA at 4°C for 5 days. The tissues were then cryoprotected in PBS containing 30% sucrose at 4°C overnight, embedded in Tissue-Tek OCT compound (Sakura Finetek), frozen on dry ice, and stored at −80°C until sectioning. Frozen samples were cut serially at 20 μm intervals with a Leica CM 3050S cryostat (Leica Biosystems Nussloch GmbH) and immunolabeled, as described below and as previously described (*47*).

Embryos and fetuses were fixed by immersion in 4% paraformaldehyde in 0.1M phosphate buffer, pH 7.4 at 4°C for 2-5 days depending on sample size. The tissues were cryoprotected in 30% sucrose/PBS at 4°C overnight, embedded in Tissue-Tek OCT compound (Sakura Finetek), frozen in dry ice and stored at −80°C until sectioning. For immunolabelling, 20 µm-thick sections of entire heads at GW 11 and GW 14 were processed as follows. Slides first underwent antigen retrieval for 20 minutes in a 5mM citrate buffer heated to 90°C, then were rinsed in TBS and blocked/permeabilized for 2 hours at room temperature in TBS + 0.3% Triton + 0.25% BSA + 5% Normal Donkey Serum (“Incubation solution”, ICS). Sections were then incubated with primary antibodies for two nights at 4°C in ICS. After rinses in TBS, the sections were incubated with secondary antibodies for two hours at RT in ICS, then rinsed again in TBS. Finally, nuclei were stained with DAPI (Sigma D9542, 1:5000 in TBS) for 5 minutes, and sections were rinsed before coverslipping with homemade Mowiol.

### Immunofluorescence labeling – Animal brains

#### Animals

##### Mice

Three C57BL/6J (Charles River) and two NPY::GFP (JAX:006417); POMC::Cre (JAX:005965); tdTomato (JAX:007914) male mice, 8–9 weeks-old were individually housed and given ad libitum access to water and standard pelleted rodent chow (R03-25, Safe diet). Three other C57BL/6J were given ad libitum water and a high-fat diet containing 60% fat (HFD; D12492 Research Diet) for 9 weeks. Ovariectomy (OVX) in mice: 6 adult female mice against a background of C57BL/6J (ERa flox with VH injection) were subjected to ovariectomy (OVX; N=3) or sham (N=3) surgery. Briefly, OVX was performed under isoflurane anesthesia. A mid-ventral incision was made, the muscle separated gently by forceps to expose ovaries and periovarian fat tissue. Ovaries and ovarian fat were removed bilaterally after ligation of the most proximal portion of the oviduct. In sham animals, the same procedure was carried out except for the removal of the ovaries. The surgical incision was sutured and postsurgical recuperation was monitored daily. Animals were kept for 6 weeks after surgery.

##### Hamsters

Two 8-week (100 g) male hamsters (Janvier) were fed ad libitum and single-housed. Animal studies were performed with the approval of the Institutional Ethics Committees for the Care and Use of Experimental Animals of the University of Lille and the French Ministry of National Education, Higher Education and Research (APAFIS#2617-2015110517317420 v5 and APAFIS#25041-2020040917227851), and under the guidelines defined by the European Union Council Directive of September 22, 2010 (2010/63/EU).

#### Brain Fixation

To fix the brains of mice and hamsters, animals were anesthetized with an intraperitoneal injection of Ketamine/Xylazine (80mg/100mg/Kg body weight). Mice were perfused transcardially with ice-cold NaCl 0.9% solution followed by the fixative solution. For wild type C57BL/6J mice fed standard chow or HFD, a solution of 4% paraformaldehyde in borate buffer (sodium tetraborate decahydrate pH 9.5) was used. For female mice, male NPY-GFP; POMC::Cre; tdTomato mice and hamsters, a fixative solution of PFA 4% in phosphate-buffered saline (PBS; pH 7.4) was used. Dissected brains were post-fixed for 4h in their respective fixative solutions before cryopreservation in sucrose 30% (sucrose in 0.1M phosphate buffered saline pH7.4) for 48h before cryosectioning. Hamsters were decapitated and the harvested brain immersion-fixed for 24h in 4% paraformaldehyde in phosphate buffer.

#### Immunohistochemistry

For triple-label immunofluorescence experiments, 30 μm-thick floating sections were rinsed 4 times in 0.1 M PBS pH 7.4 and blocked for 1 hour at room temperature in blocking solution (PBS containing 10% normal donkey serum and 0.3% Triton X-100). Sections were incubated overnight at 4°C with a mix of primary antibodies diluted in blocking solution (goat anti ACE2 1:200; rabbit anti TMPRSS2 1:1,000 and mouse anti FPR2 1:200; see Antibody table). The sections were then washed three times in 0.1M PBS and incubated for 1.5 hours at room temperature with a biotinylated donkey anti-rabbit secondary antibody to amplify the TMPRSS2 signal (1:500). The sections were then washed three times in 0.1MPBS and incubated at room temperature for 1 hour with Alexa Fluor-conjugated secondary antibodies (1:500 dilution; all purchased from Molecular Probes, Invitrogen, San Diego, CA) in blocking solution. The sections were rinsed 3 times in 0.1 M PBS. Nuclei were then counterstained by incubating the sections for 1 minute in DAPI.

### Antibody table

| *Antibody* | *Manufacturer* | *Reference* | *Dilution* | *Amplification* |
| --- | --- | --- | --- | --- |
| Goat anti human ACE2 | R&D Systems | AF933 | 1/100 | Yes |
| Goat anti mouse ACE2 | R&D Systems | AF3437 | 1/200 | No |
| Rabbit anti human ACE2 | Abcam | Ab15348 | 1/200 | No |
| Rabbit anti-TMPRSS2 | Abcam | Ab92323 | 1/100-1/1000 | Yes |
| Mouse anti-FPR2 | Invitrogen | GM1D6 | 1/100-1/400 | Yes |
| Chicken anti-vimentin | Millipore | AB5733 | 1/500 | No |
| Goat anti-TAG1 | R&D Systems | AF4439 | 1/500 | No |
| Goat anti-OMP | Wako | 544-10001 | 1/200 | No |
| Mouse anti-dsRNA | SCICONS J2 | 1001050 | 1/500 | No |
| Mouse anti-spike protein | GeneTex | GTX632604 | 1/200 | No |
| Rabbit anti-SARS nucleocapsid protein | Novus Bio | NB100-56576 | 1/100 | No |

### Fluorescence-activated cell sorting and real-time quantitative PCR

#### Tat-cre infusion

A tat-cre fusion protein produced as detailed previously (*78*) was stereotaxically infused into the third ventricle (6.37 μg/2 μl at a rate of 0.2μl/ml; AP: −1.7 mm, ML: 0 mm DV: −5.6 mm) of five isoflurane-anesthetized 4 month-old male C57Bl/6 *tdTomato*^loxP/+^ reporter mice 2 weeks before experiments, as described before (*27*).

#### Sorting

Median eminence explants were microdissected from *tdTomato*^Tat-Cre^ mice, and cell dissociated using the Papain Dissociation System (Worthington Biochemical Corporation). Fresh dissociated cells were sorted by Fluorescence-activated cell sorting (FACS) in an BD FACSAria™ III sorter.

#### Quantitative RT-PCR analyses

For gene expression analyses, lysates of FACS-sorted tanycytes and non-tanycytes were subjected to DNAse treatment (DNaseI EN0521, ThermoFisher) and then reverse transcribed using MultiScribe™ Reverse Transcriptase and the High-Capacity cDNA Reverse Transcription Kits (Applied Biosystems). A preamplification step was performed (TaqMan^tm^ PreAmp Master Mix Kit 4488593, Applied Biosystems) before real-time PCR. Real-time PCR was carried out on Applied Biosystems 7900HT Fast Real-Time PCR System using TaqMan® Gene Expression Assays listed below. Real-time PCR analysis were performed using the 2^-∆∆Ct^ method using as an internal/positive control a sample of 50pg of total RNA obtained from the mediobasal hypothalamus as extensively described previously (*79*).

### RTqPCR primer table

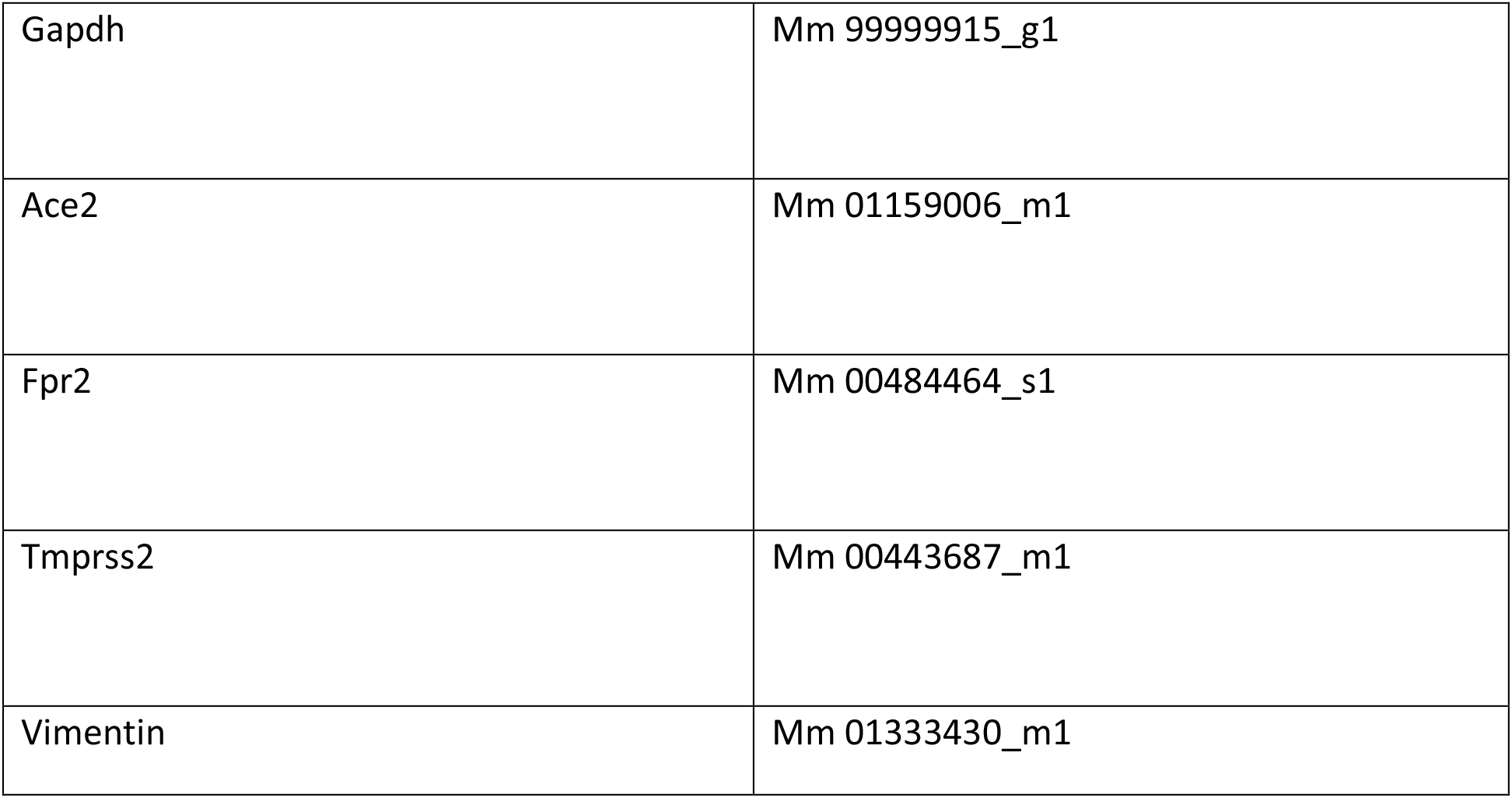

|  |  |
| --- | --- |
| Gapdh | Mm 99999915_g1 |
| Ace2 | Mm 01159006_m1 |
| Fpr2 | Mm 00484464_s1 |
| Tmprss2 | Mm 00443687_m1 |
| Vimentin | Mm 01333430_m1 |

## Supplementary Figures

**Supplementary Figure 1.**
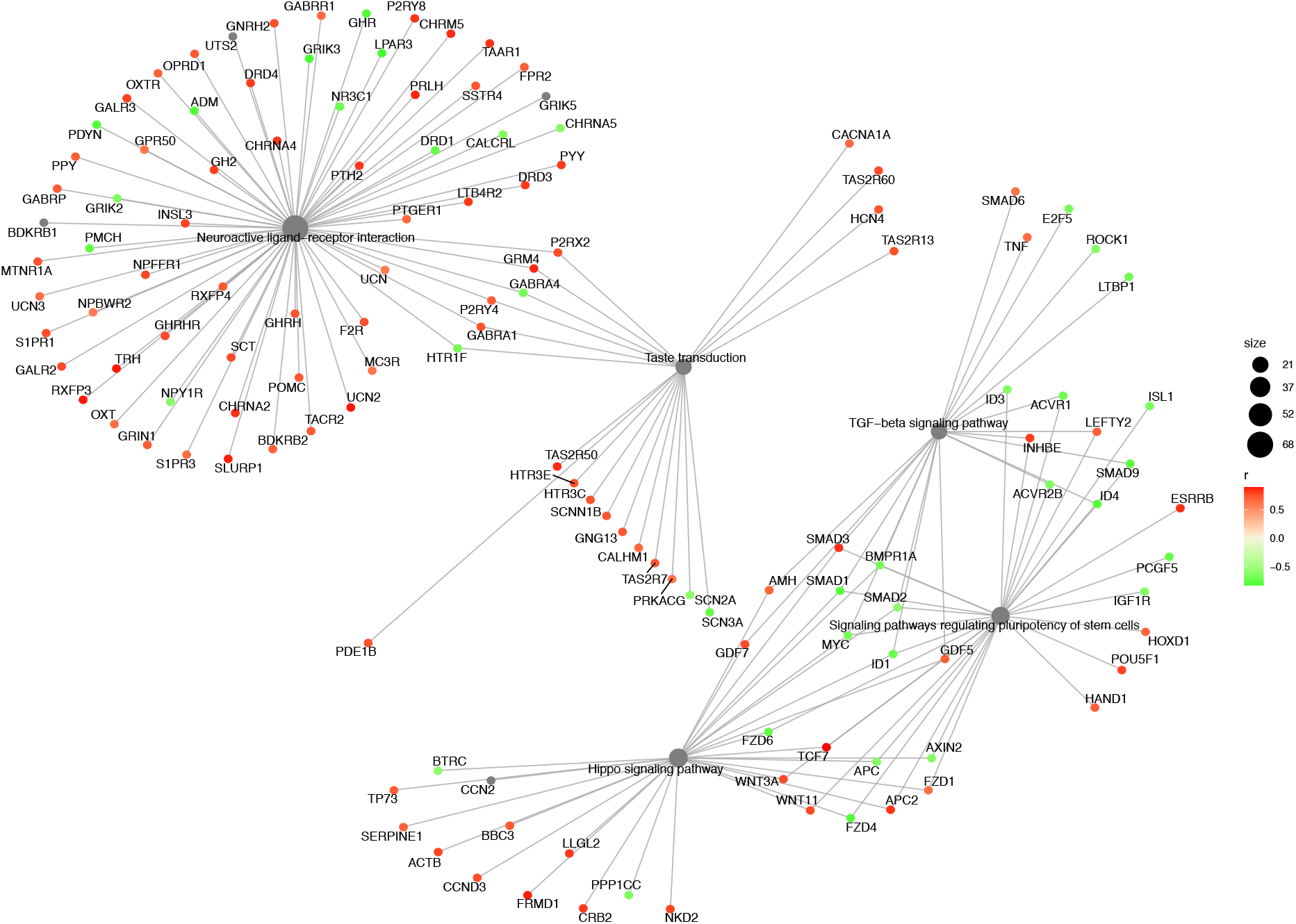
Gene network of ACE2-correlated genes enriched for KEGG pathways in the insula. Pathways enriched: “neuroactive ligand-receptor interaction”, “taste transduction”, “TGF-beta signaling”, “signaling pathways regulating pluripotency of stem cells” and “hippo signaling” pathways. Interestingly, the pathways form two subnetworks that don’t appear to interact.

**Supplementary Figure 2.**
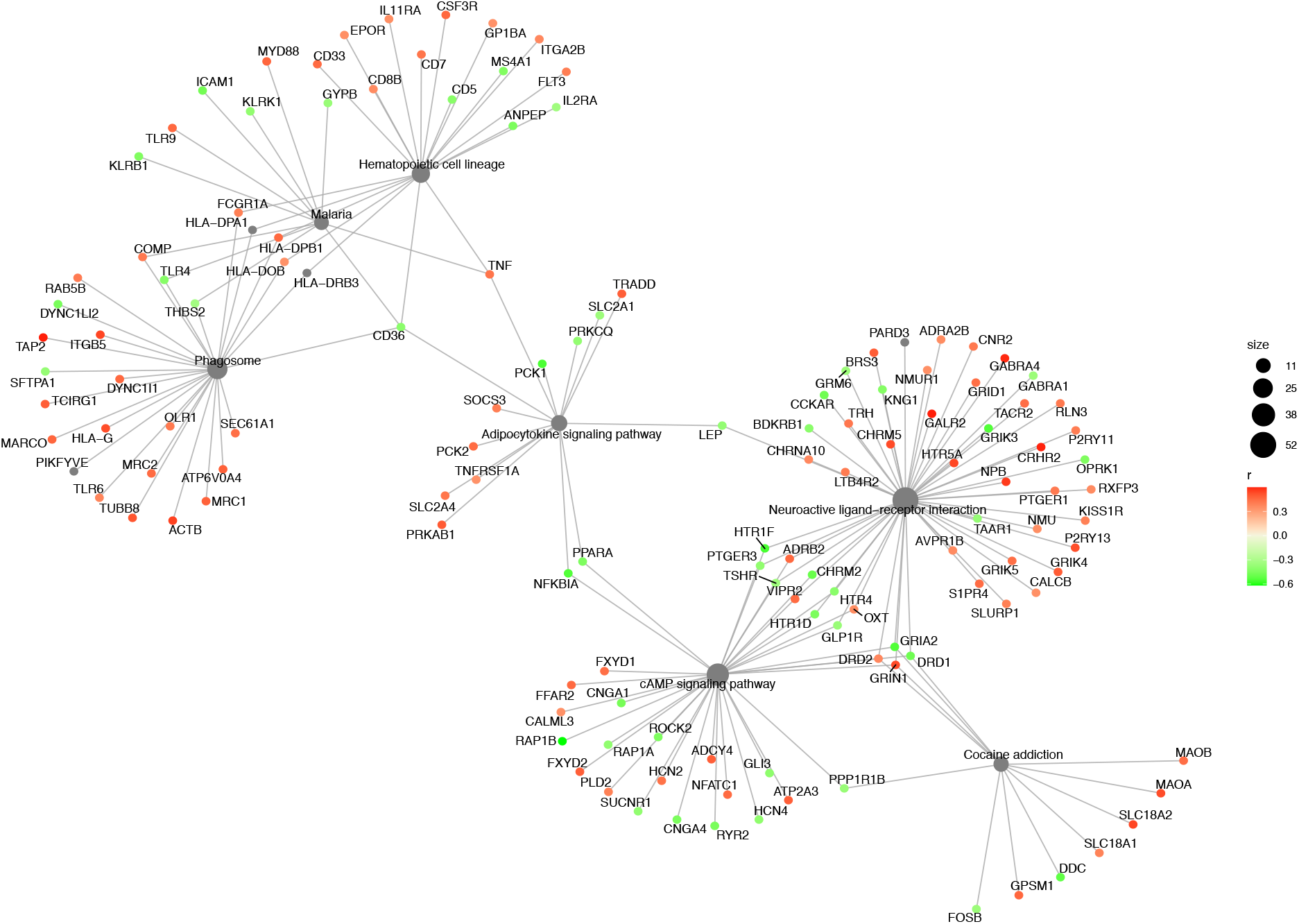
Gene network of ACE2-correlated genes enriched for KEGG pathways in the amygdala. Pathways enriched: “neuroactive ligand-receptor interaction”, “hematopoietic cell lineage”, “phagosome”, cAMP signaling”, “Cocaine addiction”, “Adipocytokine signaling”, “Malaria”. The involvement of the adipocytokine signaling pathway and the phagosome pathway might have implications for metabolic risk factors and viral entry respectively.

**Supplementary Figure 3.**
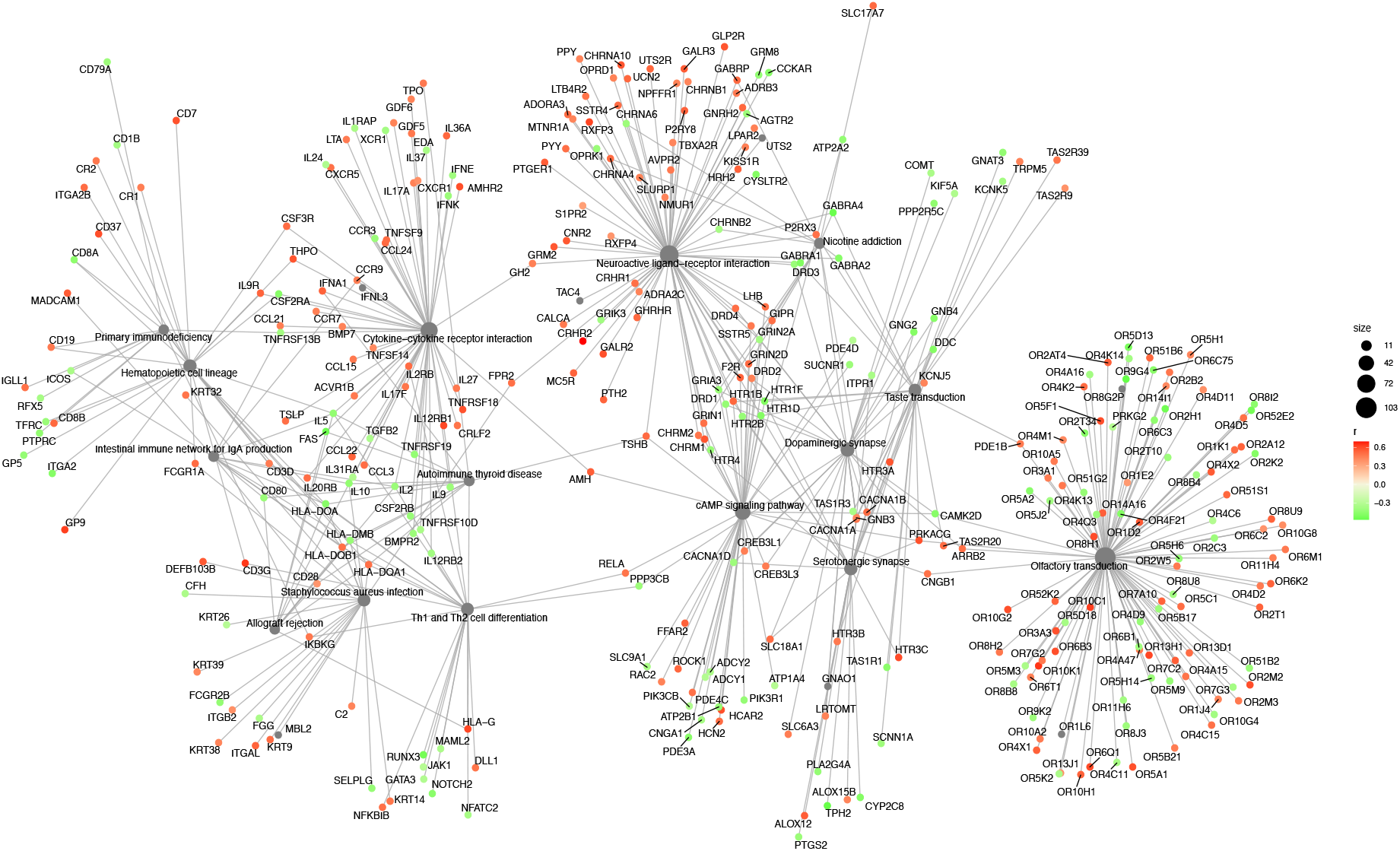
Gene network of ACE2-correlated genes enriched for KEGG pathways in the hypothalamus. Out of 45 significant pathways (fdr < 0.25), the top 15 pathways of interest have been shown in the network. “olfactory transduction”, “neuroactive ligand-receptor interaction”, “taste transduction”, “nicotine addiction”, “hematopoietic cell lineage”, “cAMP signaling pathway”, “dopaminergic synapse”, “cytokine-cytokine receptor interaction”, “Th1 and Th2 cell differentiation”, “primary immunodeficiency”, “allograft rejection”, “staphylococcus aureus infection”, “autoimmune thyroid disease”, “intestinal immune network for IgA production” pathways. The extraordinarily high number of enriched and interconnected pathways with abundant correlated genes could indicate that ACE2 plays a pivotal role across multiple hypothalamic functions, and that viral alteration of ACE2 availability or function could have wide repercussions. The occurrence of a large number of olfactory and taste receptors is intriguing, even if several of the former turn out to be pseudogenes.

**Supplementary Figure 4.**
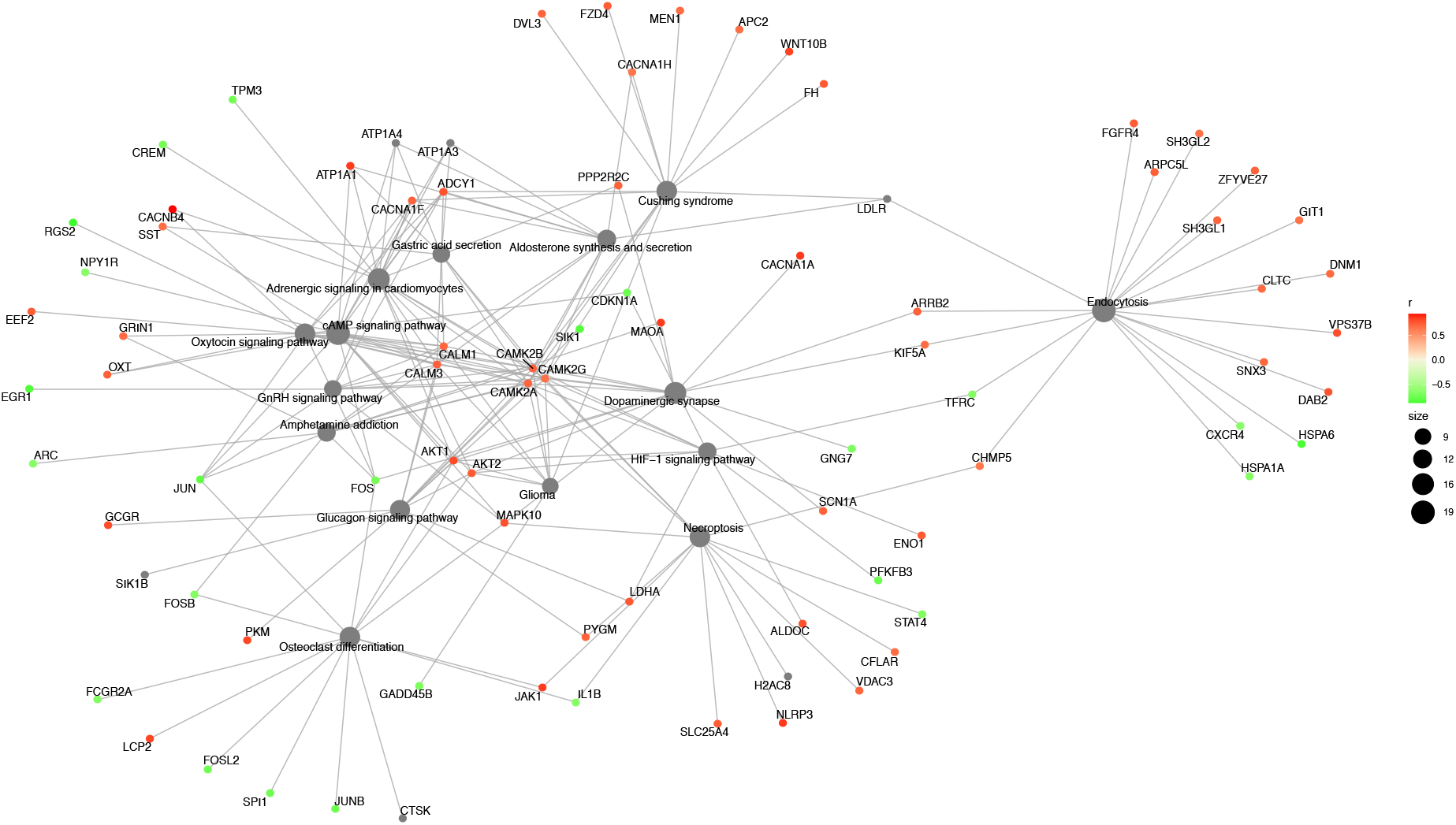
Gene network of ACE2-correlated genes enriched for KEGG pathways in the parabrachial Nuclei of Pons. Out of 73 significant pathways (fdr < 0.25), the top 15 pathways of interest have been shown in the network. Pathways enriched: “dopaminergic synapse”, “amphetamine addiction”, “adrenergic signaling in cardiomyocytes”, “glucagon signaling pathway”, “aldosterone synthesis and secretion”, “osteoclast differentiation”, “cAMP signaling”, “gastric acid secretion”, “glioma”, “oxytocin signaling pathway”, “Cushing syndrome”, “endocytosis”, “GnRH signaling”, “HIF-1 signaling”, “necroptosis” pathways. In contrast to the hypothalamus, however, the number of genes per enriched pathway is fairly low, even though some genes, calmodulins and calmodulin kinases, appear to be common to a very large number of them.

**Supplementary Figure 5.**
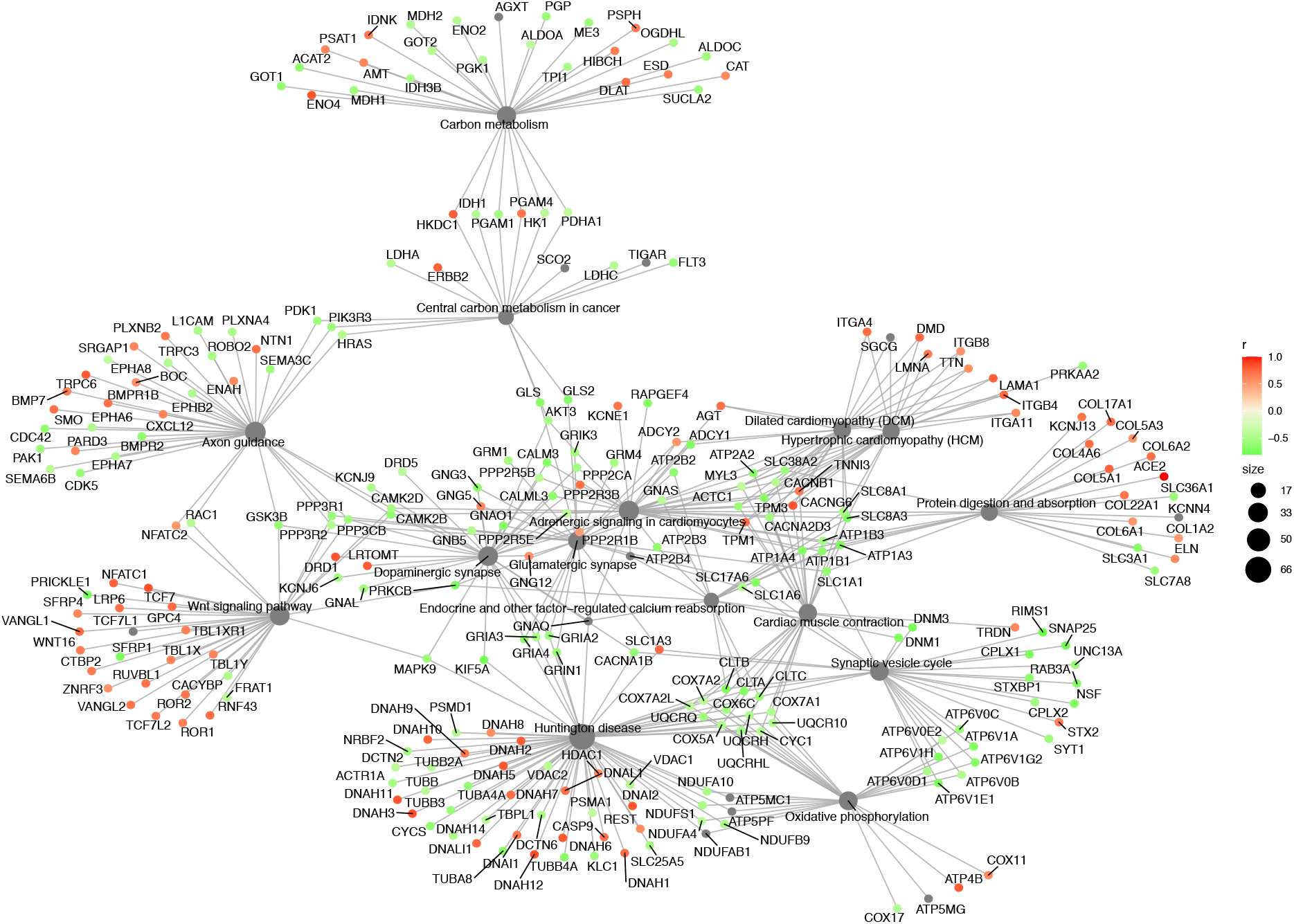
Gene network of ACE2-correlated genes enriched for KEGG pathways in the myelencephalon. Out of 73 significant pathways (fdr < 0.25), the top 15 pathways of interest have been shown in the network “Huntington disease”, “synaptic vesicle cycle”, “cardiac muscle contraction”, “carbon metabolism”, “adrenergic signaling in cardiomyocytes”, “dopaminergic synapse”, “endocrine and other factor-regulated calcium reabsorption”, “glutamatergic synapse”, “dilated cardiomyopathy”, “protein digestion and absorption”, “oxidative phosphorylation”, “hypertrophic cardiomyopathy (HCM)”, “Wnt signaling pathway”, “axon guidance”, “central carbon metabolism in cancer” pathways. The myelencephalon, like the hypothalamus has a very large number of networks with abundant ACE2-correlated genes, in keeping with the wide range of functions that are coordinated in this region. This could contribute to the potential vulnerability of cardiorespiratory centers to infection.

**Supplementary Figure 6.**
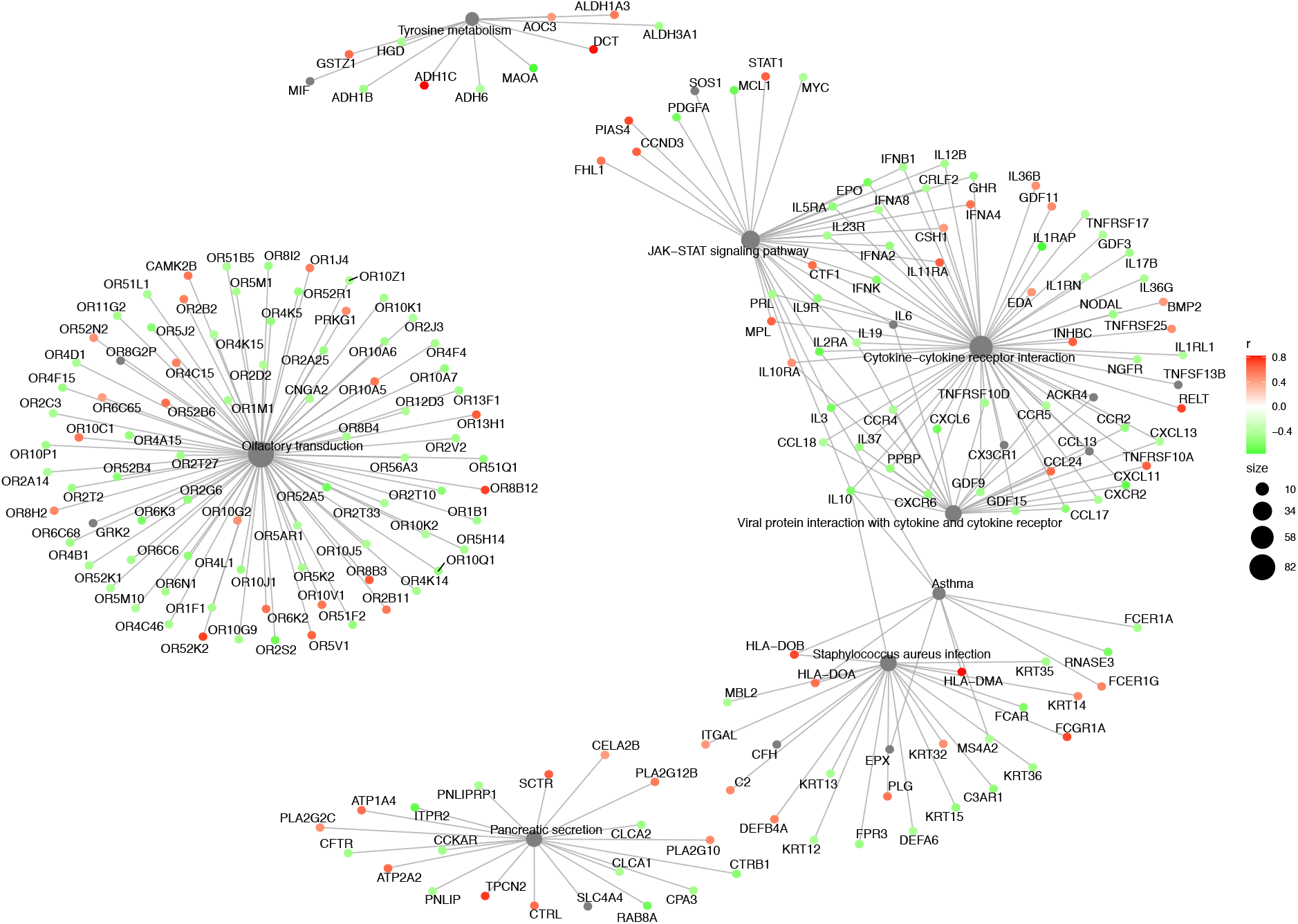
Gene network of TMPRSS2-correlated genes enriched for KEGG pathways in the insula. Pathways enriched: “olfactory transduction”, “tyrosine metabolism”, “JAK-STAT signaling”, cytokine-cytokine signaling pathway, asthma, “Staphylococcus aureus infection”, “pancreatic secretion” and “viral protein interaction with cytokine and cytokine receptor” pathways. Similar to the network for ACE2-correlated genes in the insula, the TMPRSS2-correlated genes also form several subnetworks that do not interact much. Here again, the abundance of OR genes is intriguing, as is the apparent enrichment for infectious and inflammatory pathways.

**Supplementary Figure 7.**
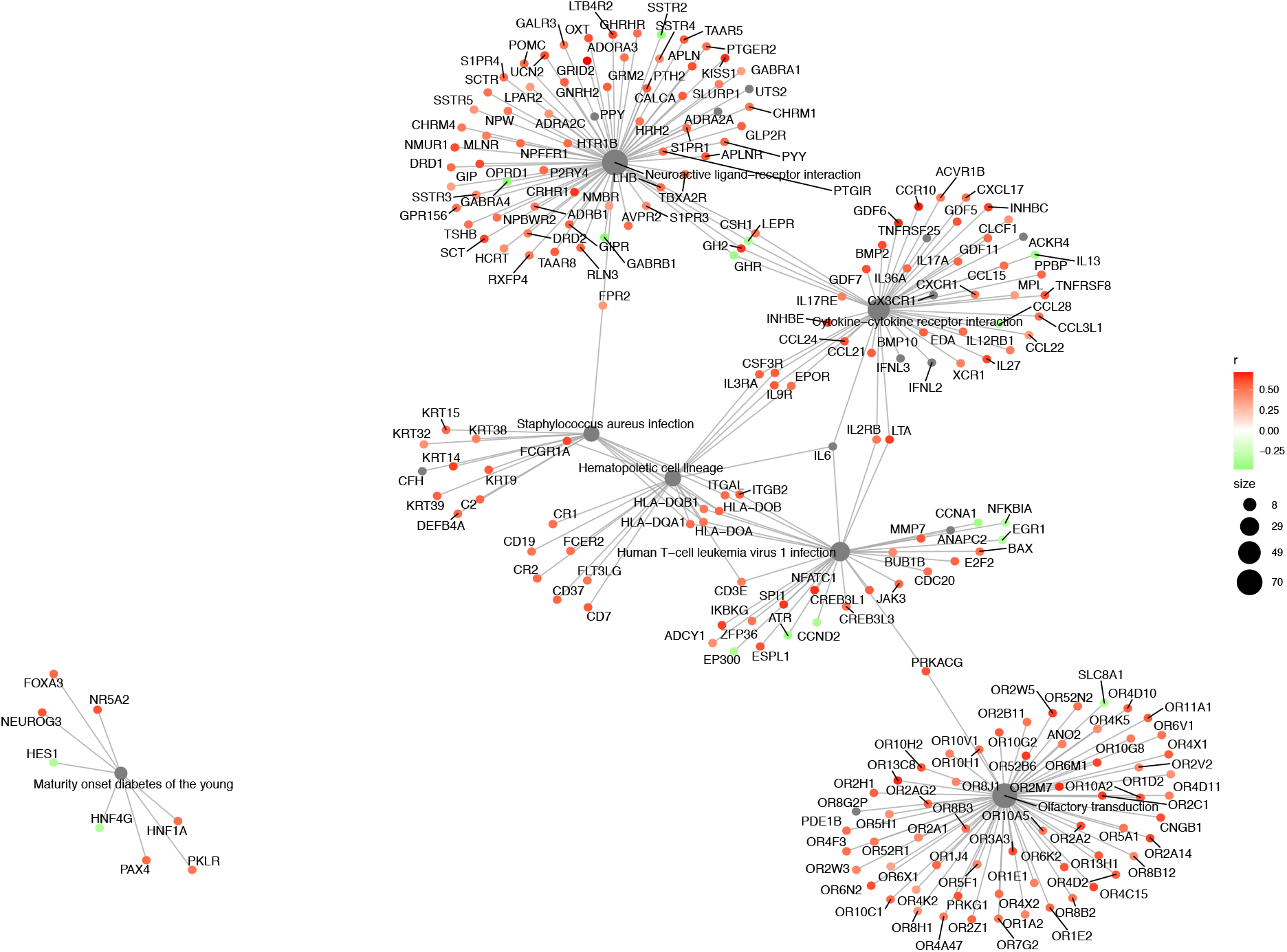
Gene network of TMPRSS2-correlated genes enriched for KEGG pathways in the amygdala. Pathways enriched: “neuroactive ligand-receptor interaction”, “olfactory transduction”, “cytokine-cytokine receptor interaction”, “maturity onset of diabetes in the young”, “hematopoietic cell lineage”, “human T-cell leukemia virus 1 infection”, and “Staphylococcus aureus infection” pathways. As in the insula, there is a pattern of inflammatory or infectious pathways connecting other pathways.

**Supplementary Figure 8.**
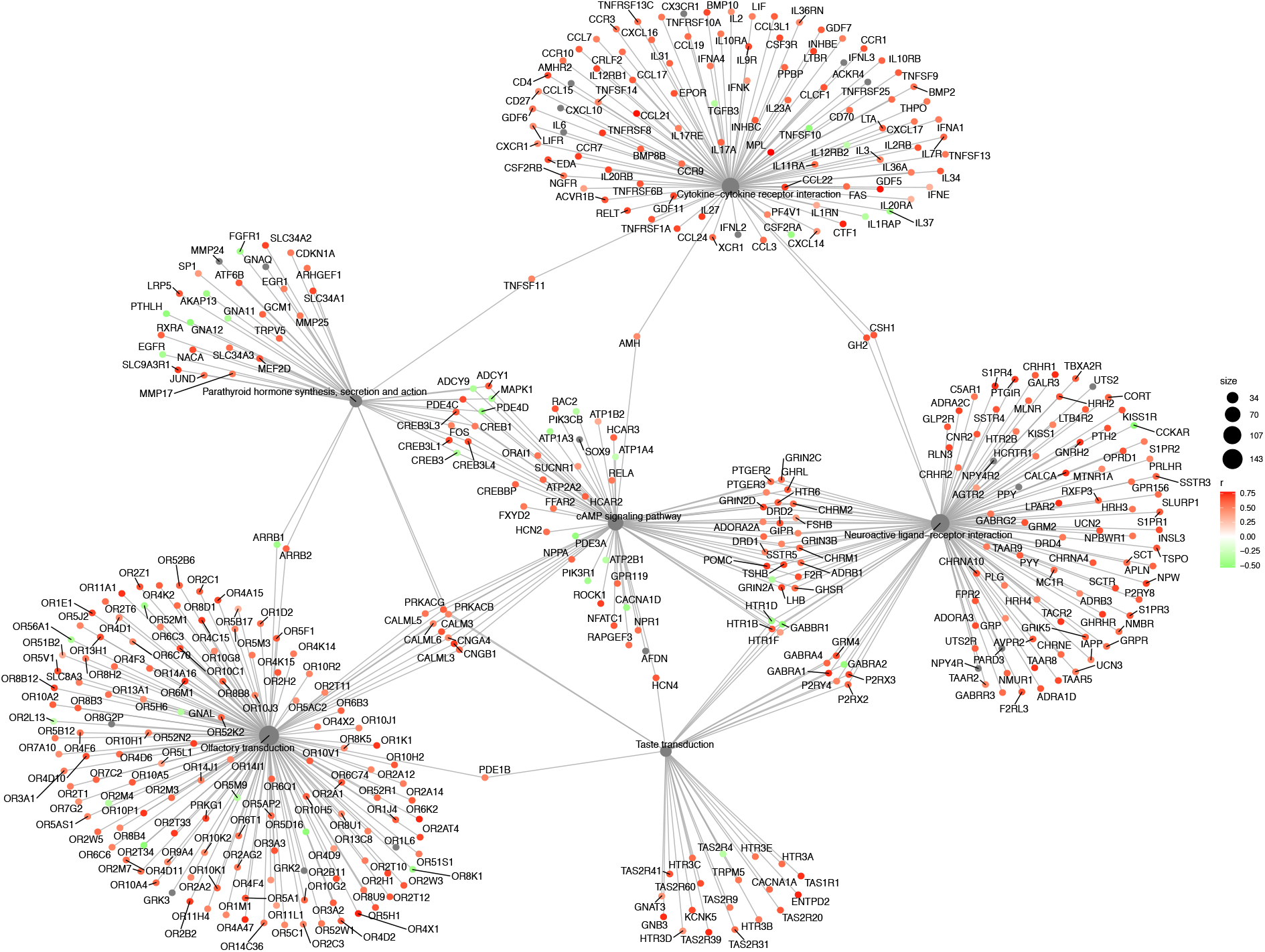
Gene network of TMPRSS2-correlated genes enriched for KEGG pathways in the hypothalamus. Pathways enriched: “neuroactive ligand-receptor interaction”, “olfactory transduction”, “parathyroid hormone synthesis, secretion and action”, “taste transduction”, cAMP signaling”, pathways. The hypothalamic network for TMPRSS2-enriched genes appears less dense than that for ACE2-correlated genes. cAMP signaling appears to be a common point of interaction for diverse other networks, which do not interact much between themselves, with one or two exceptions.

**Supplementary Figure 9.**
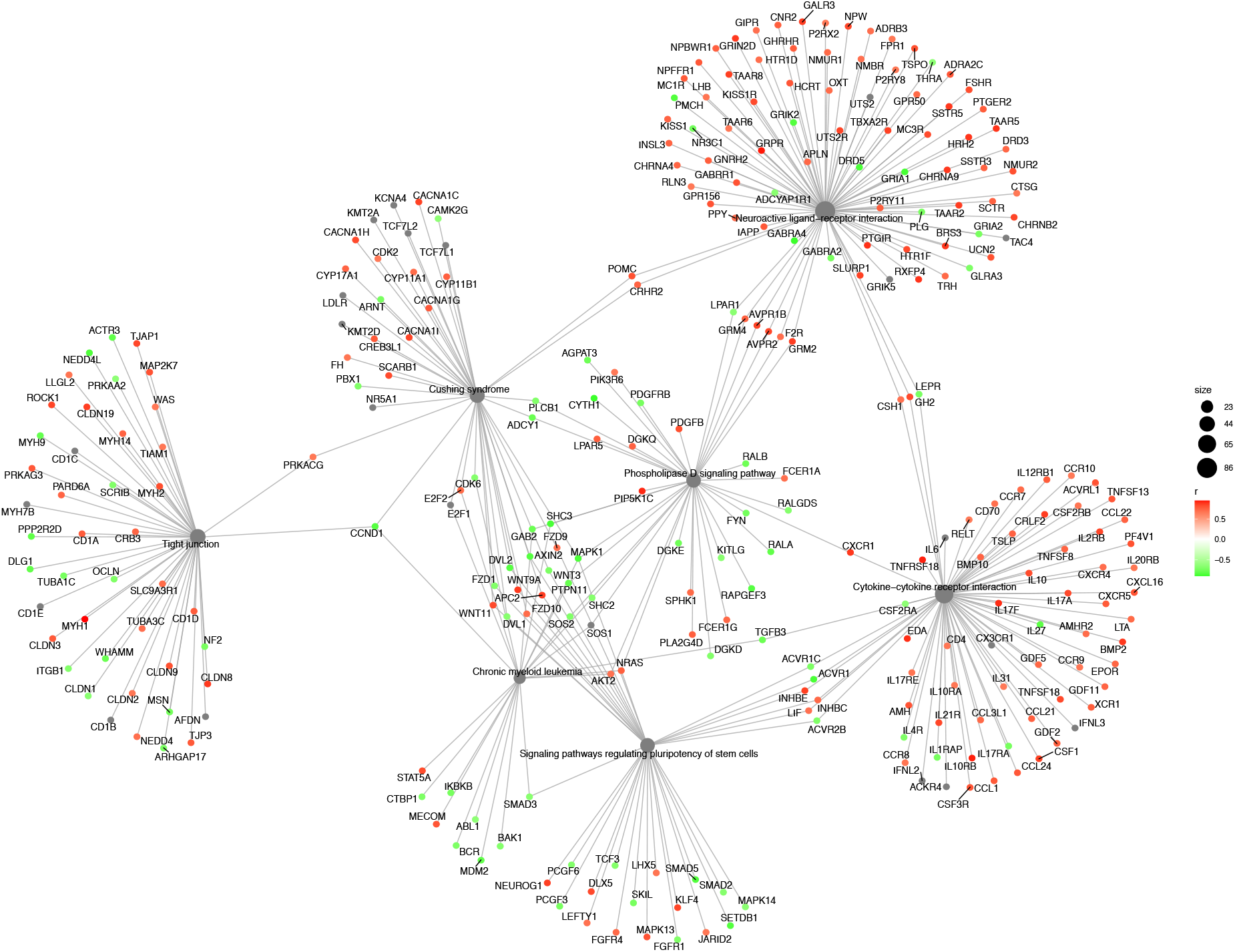
Gene network of TMPRSS2-correlated genes enriched for KEGG pathways in the parabrachial nuclei of pons. Pathways enriched: “neuroactive ligand-receptor interaction”, “tight junction”, “Cushing syndrome”, “cytokine-cytokine receptor interaction”, “chronic myeloid leukemia”, “signaling pathways regulating pluripotency of stem cells”, “phospholipase D signaling” pathways.

**Supplementary Figure 10.**
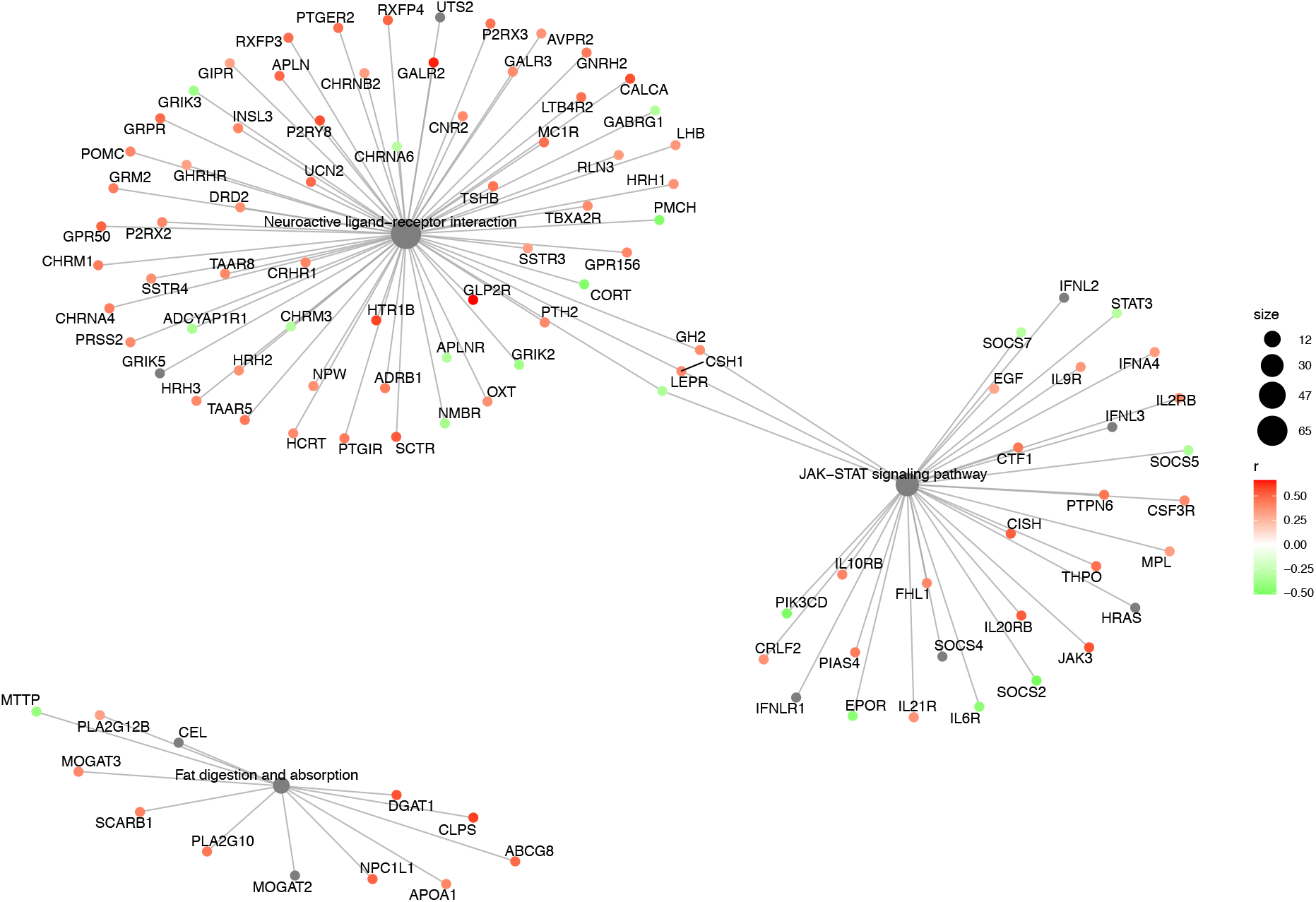
Gene network of TMPRSS2-correlated genes enriched for KEGG pathways in the myelencephalon. Pathways enriched: “neuroactive ligand-receptor interaction”, “JAK-STAT signaling pathway”, “fat digestion and absorption”. Contrary to the diversity of enriched pathways for ACE2-correlated genes, only 3 pathways were yielded for TMPRSS2-correlated genes.

**Supplementary Figure 11.**
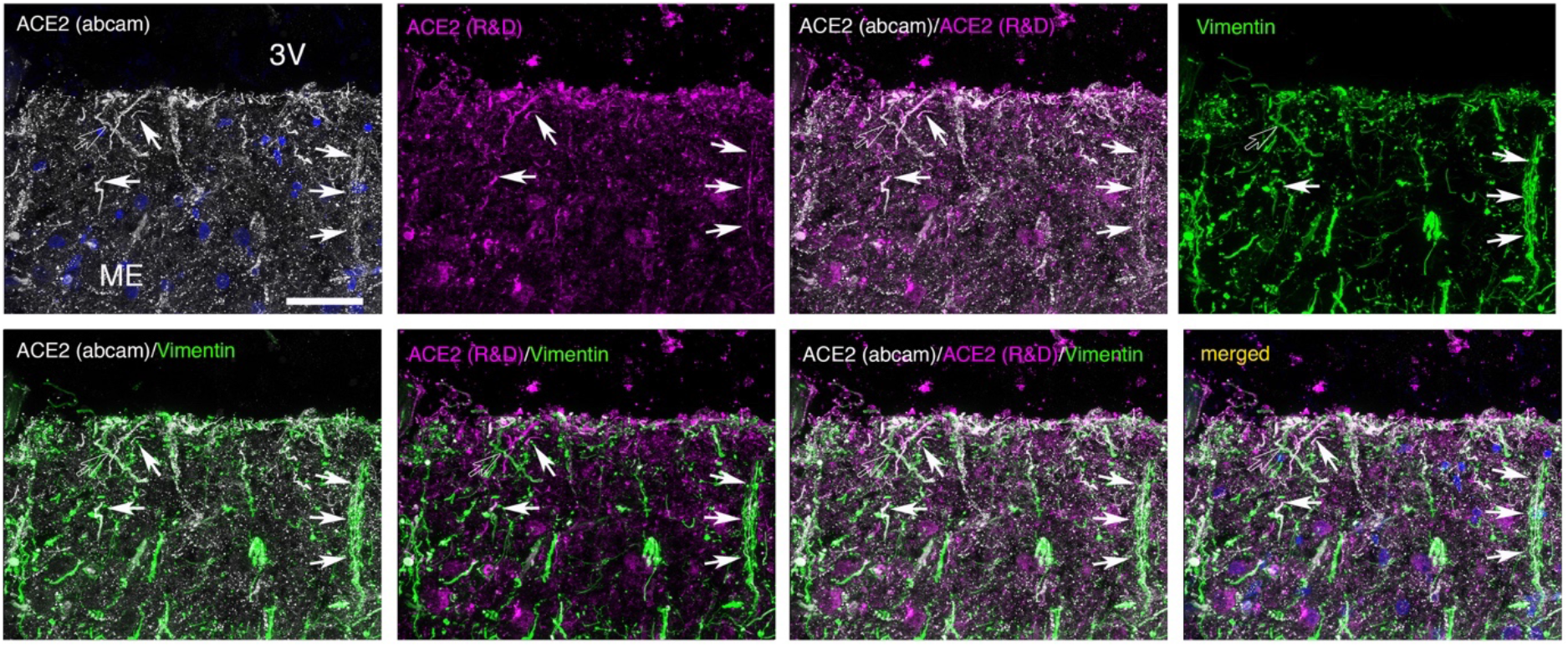
Similar immunofluorescence patterns obtained with two antibodies to human ACE2 in the median eminence (ME) of the 63 year-old COVID-19 patient. Colabeling with vimentin indicates that the fibers expressing ACE2 in the two cases belong to tanycytes. Scale bars: 40μm.

## Supplementary Excel. Tables

**Supplementary Table 1. log2 expression values for ACE2 and TMPRSS2 across selected brain regions in the AHBA.** Expression levels (median log2 intensity)of ACE2 and TMPRSS2 in all the brain regions and reference structures selected are listed here.

**Supplementary Table 2. List of all genes correlated with ACE2 and/or TMPRSS2 in the 5 regions analyzed.** Pearson’s rank correlation coefficient (r), p value and adjusted p value for all the genes found to be significantly correlated with either ACE2 or TMPRSS2 in each of the 5 regions.

**Supplementary Table 3. List of genes correlated with ACE2 and/or TMPRSS2 in the hypothalamus and common to the COVID-19 lung geneset.** Genes significantly correlated with either ACE2 or TMPRSS2 or both in the hypothalamus, and also present in the geneset of differentially expressed genes from the COVID-19 lung (*20*).

**Supplementary Table 4. GO enrichment terms of ACE2-correlated genes.** In addition to KEGG pathways, we performed enrichment for GO terms using our ACE2-correlated genes in the 5 brain regions of interest. The list of GO terms yielded for “biological processes”, “cellular components” and “molecular function” are presented.

**Supplementary Table 5. GO enrichment terms of TMPRSS2-correlated genes.** In addition to KEGG pathways, we performed enrichment for GO terms using our TMPRSS2-correlated genes in the 5 brain regions of interest. The list of GO terms yielded for “biological processes”, “cellular components” and “molecular function” are presented.

### Supplementary Movie

**Movie1. Representation of ACE2 and TMPRSS2 gene expression levels in brain regions of interest in the AHBA.**

